# A single C-terminal residue controls SARS-CoV-2 spike trafficking and virion assembly

**DOI:** 10.1101/2023.03.09.531992

**Authors:** Debajit Dey, Enya Qing, Yanan He, Yihong Chen, Benjamin Jennings, Whitaker Cohn, Suruchi Singh, Lokesh Gakhar, Nicholas J. Schnicker, Brian G. Pierce, Julian P. Whitelegge, Balraj Doray, John P. Orban, Tom Gallagher, S. Saif Hasan

## Abstract

The spike (S) protein of SARS-CoV-2 is delivered to the virion assembly site in the ER-Golgi Intermediate Compartment (ERGIC) from both the ER and cis-Golgi in infected cells^1-3^. However, the relevance and modulatory mechanism of this bidirectional trafficking are unclear. Here, using structure-function analyses, we show that S incorporation into virions and viral fusogenicity are determined by coatomer-dependent S delivery from the cis-Golgi and restricted by S-coatomer dissociation. Although S mimicry of the host coatomer-binding dibasic motif ensures retrograde trafficking to the ERGIC, avoidance of the host-like C-terminal acidic residue is critical for S-coatomer dissociation and therefore incorporation into virions or export for cell-cell fusion. Because this C-terminal residue is the key determinant of SARS-CoV-2 assembly and fusogenicity, our work provides a framework for the export of S protein encoded in genetic vaccines for surface display and immune activation.

## INTRODUCTION

COVID-19 is caused by the coronavirus SARS-CoV-2, the most thoroughly studied virus to date, which is targeted by multiple different vaccines^4^. The SARS-CoV-2 virion comprises genomic RNA complexed with cytoplasmic nucleocapsid (N), which is surrounded by a lipid membrane containing S, membrane (M), and envelope (E) proteins^1,5-8^. Following infection of host cells, SARS-CoV-2 replicates and traffics its components to the ERGIC, where virions are assembled for subsequent release from the cell. Viral assembly can occur in the absence of the S protein, but incorporation of S is required for the virion to bind, enter, and infect new host cells^9-11^.

During normal cellular secretion, anterograde trafficking exports endogenous cargo from the ER whereas retrograde trafficking retrieves proteins such as type I membrane proteins that escape from the ER during anterograde trafficking^12-17^. These escaped client proteins are recognized in the cis-Golgi network by cytosolic coatomer protein I (COPI), a 560 kDa complex of seven distinct gene products (α-, β-, β′-, γ-, δ-, ε-, and ζ-COPI), which assemble to form the coatomer^18-24^ **(Figure 1A)**. The coatomer binds to a C-terminal dibasic retrieval motif, Lys-x-Lys-x-x_CT_ or Lys-Lys-x-x_CT_ (x = any amino acid; x_CT_ = any C-terminal amino acid), which leads to client packaging into COPI-coated vesicles and retrieval to ER^25-31^. The N-terminal β-propellor WD40 domains of α and β’ coatomer subunits (αWD40 and β’WD40, respectively) provide binding sites for the dibasic motif on client proteins^28,32-34^. SARS-CoV-2 hijacks this secretory system to assemble its components into virions. In particular, the S protein – a type I membrane protein – is delivered to the ERGIC from both the ER (by anterograde trafficking) and the Golgi (by retrograde trafficking)^1,2,35,36^ **(Figure 1A)**. Retrograde delivery of S is due to a Lys-x-His-x-x_CT_ sequence in its cytosolic tail, which partially mimics the coatomer-binding dibasic motif^2,35^. However, x_CT_ is frequently acidic in endogenous client proteins and strictly non-acidic in CoV spike molecules^34^, revealing an abrupt departure from mimicry immediately downstream of the dibasic motif (S protein extended motif). The significance of this bipartite mimicry and how it modulates incorporation of the S protein into virions is unclear. Moreover, little is known about the relevance of bidirectional S trafficking during virion assembly.

**Figure 1.**
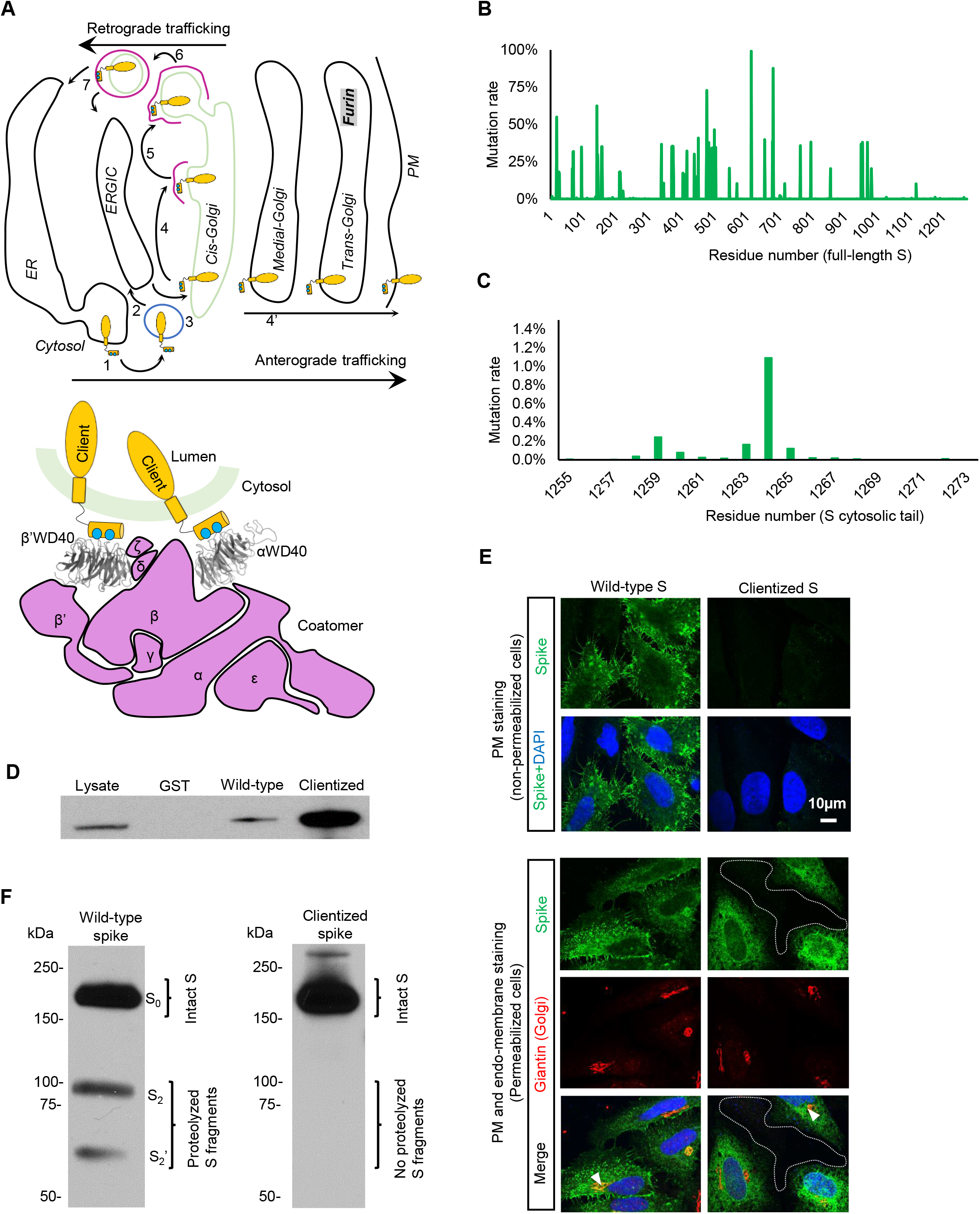
The conserved C-terminus determines S protein trafficking and coatomer interactions. **(A)** Left panel: Schematic of bidirectional trafficking of S in the secretory pathway. The newly synthesized S protein (yellow) in the ER (1) is exported in vesicles (blue), trafficked to the ERGIC (2), and delivered to the cis-Golgi (3). COPI-coated vesicles (pink) retrieve S to the ER and ERGIC (4-7) by retrograde trafficking. S not retrieved by COPI is exported to the PM (4’) via the furin-containing trans-Golgi by anterograde trafficking. Right panel: The coatomer is a hetero-heptamer of seven distinct gene products. WD40 domains on α and β’ subunits form the binding site for client proteins such as S. **(B)** Multiple mutational hotspots in the N-terminal two-thirds of the full-length SARS-CoV-2 S, which corresponds to the ectodomain. **(C)** Few mutations occur in the S tail. **(D)** Clientized S tail-GST fusion protein pulls down ∼ 17-fold more coatomer than wild-type S tail-GST fusion. **(E)** Immunofluorescence imaging of full-length wild-type and clientized S in permeabilized and non-permeabilized HeLa cells. Wild-type S localizes to the PM and early secretory compartments. Clientized S protein is absent from the PM and restricted to early secretory compartments. White outline, PM; white arrowheads, early secretory compartments. **(F)** Western blots of HeLa cells expressing wild-type or clientized full-length S. Wild-type S yields intact, unproteolyzed fraction from early secretory compartments and proteolyzed, furin-cleaved fragments from PM (left). Clientized S yields only an intact fraction from early secretory compartments (right).S_0_, intact S; S_1_, S_2_, S_2_’, proteolyzed fragments.

Here we utilize recombinant SARS-CoV-2 S proteins and tail peptides, including a construct with enhanced mimicry of the coatomer binding motif (clientized S protein), to elucidate the atomic structure of the S-coatomer interface. We also provide functional insight into the unconventional single-residue modification of the C-terminal retrieval motif, revealing its role in S-coatomer dissociation and incorporation of S into virions. Finally, we reveal the importance of retrograde trafficking for fusogenicity and propagation of SARS-CoV-2.

## RESULTS

### The conserved C-terminus determines S protein trafficking and coatomer interactions

To determine whether residues in the SARS-CoV-2 S tail are under selection pressure, we analyzed over 11 million sequences of full-length SARS-CoV-2 S in the GISAID database^37,38^ **(Figure 1B)**. Residues predicted to make substantial contact with the coatomer surface (Lys1269, His1271, and Thr1273) showed minimal mutation rates (0.0020%, 0.0015%, and 0.0004%, respectively) **(Figure 1C)**. The highest frequency mutation at the C-terminus was Thr1273Ser. Importantly, mutations predicted to increase mimicry by the S protein extended motif (Thr1273Asp or Thr1273Glu), were not reported.

We subsequently asked whether such clientizing mutations modify coatomer interactions *in vivo* and localization in secretory compartments using a pull-down assay of the endogenous coatomer complex with an N-terminal glutathione S-transferase (GST) fusion of the S tail (Leu1244 to Thr1273; **Figure 1D; Extended Data Figure S1)**. Clientized GST-S tail constructs containing either Thr1273Asp or Thr1273Glu showed 17-fold higher affinity for the β-COPI subunit (a marker for the intact coatomer complex) than the wild-type GST-S tail construct **(Figure 1D; Extended Data Figure S1)**. Furthermore, these clientizing mutations enhanced binding by 2-fold compared to the canonical S tail <u>Lys</u>-x-<u>Lys</u>-x-x_CT_ dibasic mutant. Mass spectrometry identified all seven coatomer subunits in pull-downs with GST-S tail clientized constructs **(Extended Data Table S1)**. These data show that clientizing mutations outside the dibasic motif enhance coatomer binding affinity compared to the endogenous canonical <u>Lys</u>-x-<u>Lys</u>-x-x_CT_ motif, suggesting that the S x_CT_ residue has a key modulatory function in coatomer interactions.

Given that the coatomer mainly associates with early secretory compartments, we evaluated whether longer-lived coatomer association due to clientizing mutations alters the localization of S proteins. Immunofluorescence microscopy showed full-length wild-type S both at the PM and in secretory compartments **(Figure 1E)**. In contrast, clientized S carrying the Thr1273Glu mutation became sequestered in coatomer-enriched early secretory compartments and did not reach the PM **(Figure 1E)**. We next asked whether relocalization of clientized S was due to lack of export from early secretory compartments or recycling from the PM **(Figure 1A)**. Cleavage of S protein into S1 and S2 fragments by the trans-Golgi protease furin is a marker for trafficking out of early secretory compartments^39-42^ **(Figure 1A)**. We observed both an intact band and proteolyzed fragments for wild-type S, corresponding to secretory compartment- and PM-localized populations, respectively **(Figure 1F)**. In contrast, a single, intact band was observed for clientized S, corresponding to retention in early secretory compartments rather than recycling from the PM **(Figure 1F)**. Together, these data demonstrate that the C-terminus, immediately downstream of the dibasic motif, is a critical modulator of S trafficking.

### The clientized S protein C-terminus engages a basic cluster in WD40 domains

The αWD40 and β’WD40 domains are structural homologs with complete conservation of the binding site except for two residues^43,44^. The SARS-CoV-2 tail displays selectivity for αWD40 over β’WD40^34^. To determine an atomic-level understanding of S-coatomer interactions, we attempted co-crystallization of αWD40 with either wild-type (1267Gly-Val-<u>Lys</u>-Leu-<u>His</u>-Tyr-Thr1273) or clientized (1267Gly-Val-<u>Lys</u>-Leu-<u>His</u>-Tyr-Glu1273) S tail heptapeptides. However, contacts between symmetry-related αWD40 molecules occluded the binding site in these crystals and no peptide density was observed. We subsequently adopted a previously described strategy of using β’WD40 for co-crystallization of peptides containing the Lys-x-His-x-x_CT_ motif^44^. The β’WD40 co-crystal structures with the wild-type and clientized S heptapeptides were determined to a resolution of 1.4 Å **(Extended Data Table S2)** and reveal substantial similarity (Cα RMSD=0.2Å). The major difference is that whereas only the main chain carboxylate of wild-type Thr1273 interacts with basic Lys17 on the β’WD40 surface, both the side chain and main chain carboxylate groups of the clientized Glu1273 residue form extensive interactions with a β’WD40 basic cluster of Arg15, Lys17, and Arg272 **(Figure 2A, B)**. This suggests that interactions between Glu1273 and the basic cluster underlie the enhanced binding of the clientized S protein. Overall, the two heptapeptides occupy the canonical WD40 binding site, which provides electrostatic complementarity to the dibasic motif residues, Lys1269 and His1271 **(Figure 2C; Extended Data Figure S2A, B)**.

**Figure 2.**
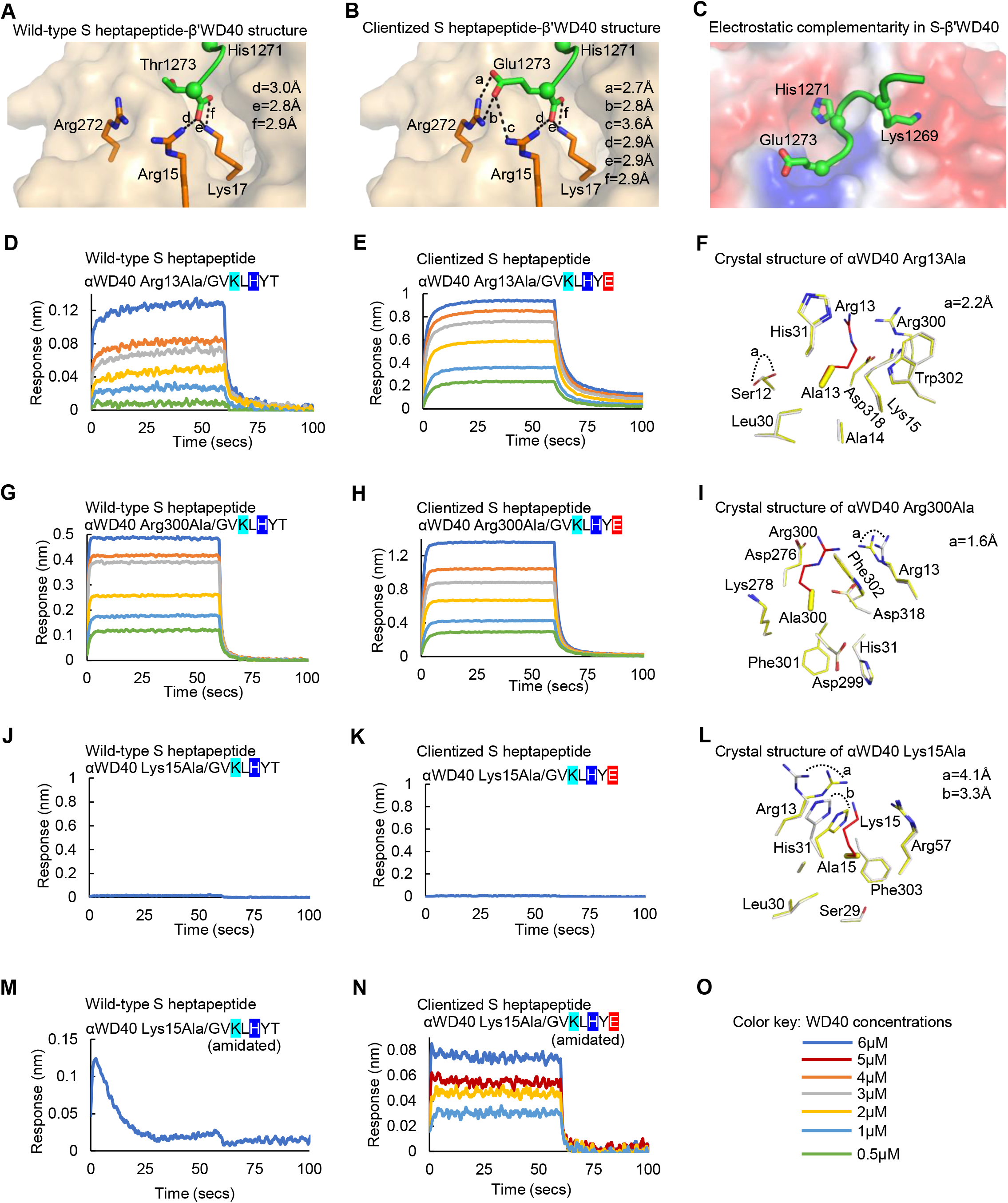
Clientized S binds to a basic cluster on the coatomer WD40 domain. **(A-C)** Co-crystal structures of SARS-CoV-2 S tail heptapeptides (residues 1267-1273) in complex with β’WD40. **(A)** Glu1273 in clientized S forms complementary electrostatic interactions with the β’WD40 basic cluster. **(B)** Thr1273 in wild-type S shows no interactions with the basic cluster. **(C)** The β’WD40 surface provides electrostatic complementarity to the S dibasic motif and the C-terminus. Color code: red, acidic; blue, basic. **(D, E)** BLI assays for αWD40 Arg13Ala mutant and wild-type S heptapeptide **(D)** or Thr1273Glu clientized S heptapeptide **(E)** show Arg13Ala has weaker binding affinity for wild-type than clientized S. **(F)** Superposition of wild-type (white) and Arg13Ala mutant (yellow) crystal structures shows minimal perturbation near the mutation site. **(G, H)** BLI assays for αWD40 Arg300Ala mutant and wild-type S heptapeptide **(G)** or Thr1273Glu clientized S heptapeptide **(H)** show Arg300Ala abolishes enhancement from clientization and has similar affinity for wild-type and clientized S. **(I)** Superposition of wild-type (white) and Arg300Ala mutant (yellow) crystal structures shows minimal perturbation near the mutation site. **(J, K)** BLI assays for αWD40 Lys15Ala mutant and wild-type S heptapeptide **(J)** or Thr1273Glu clientized S heptapeptide **(K)** show Lys15Ala abolishes binding to wild-type and clientized S heptapeptide. **(L)** Superposition of wild-type (white) and Lys15Ala mutant (yellow) crystal structures shows substantial perturbation and inward rotation of Arg13 and His31 side chains to block the S heptapeptide binding site. **(M, N)** BLI assays for amidated αWD40 Lys15Ala mutant and wild-type S heptapeptide **(M)** or Thr1273Glu clientized S heptapeptide **(N)** show amidation of the C-terminal main chain carboxylate abolishes αWD40 binding to wild-type but not clientized S heptapeptide. **(O)** Color key indicates αWD40 concentrations for BLI assays.

To gain insight into the role of the basic cluster in stabilizing S-coatomer interactions, we generated alanine mutants of αWD40 Arg13, Lys15, and Arg300, which are equivalent to the β’WD40 basic cluster residues Arg15, Lys17, and Arg272, respectively. Biolayer interferometry (BLI) assays showed that Arg13Ala weakened αWD40 binding to both wild-type and clientized S heptapeptides, although the affinity for the clientized S heptapeptide was still 2.8-fold higher **(Table 1, Figure 2D, E)**. A 1.9 Å resolution crystal structure of this αWD40 Arg13Ala mutant showed a binding site architecture comparable to that in wild-type αWD40 **(Figure 2F; Extended Data Figure S2C; Extended Data Table S2)**, attributing affinity weakening to the alanine mutation. The Arg300Ala mutation yielded similar dissociation constants (K_D_s) of 3.6 ± 0.4 μM and 4.2 ± 0.4 μM for wild-type and clientized S heptapeptides, respectively **(Figure 2G-I; Extended Data Table S2)**, and a 1.8 Å resolution crystal structure showed minimal perturbation of the binding site **(Figure 2I; Extended Data Figure S2D)**. This suggests that the Arg300 side chain is important for the higher affinity for the clientized S heptapeptide. Finally, the Lys15Ala mutant did not bind to either the wild-type or clientized S heptapeptides **(Figure 2J, K)**, and a 1.6 Å resolution crystal structure revealed this was due to conformational occlusion of the αWD40 binding site by an inward rotation of nearby Arg13 and His31 side chains, rather than loss of Lys15 interactions **(Figure 2L; Extended Data Figure S2E; Extended Data Table S2)**. We therefore generated wild-type and clientized S heptapeptides with amidation at the C-terminal main chain carboxylate, which neutralizes the main chain charge that interacts with αWD40 Lys15 without perturbing the binding site architecture of αWD40. Loss of αWD40 binding was observed for the amidated wild-type S heptapeptide **(Figure 2M, Table 1)** but not the amidated clientized S heptapeptide **(Figure 2N, Table 1)**. Hence, the Glu1273 side chain in the clientized S protein provides partial compensation for the loss of main chain stabilization from the WD40 binding site, confirming the importance of this acidic residue for WD40 domain binding.

**Table 1:** Dissociation and rate constants for the interaction of spike tail peptides with coatomer WD40 domains.

| Peptide sequence<sup>1</sup> | WD40 analyte | Equilibrium KD (μM) | k<sub>on</sub> (10<sup>3</sup>/Ms) | k<sub>off</sub> (10<sup>-1</sup>/s) | χ<sup>2</sup> |
| --- | --- | --- | --- | --- | --- |
| GVKLHAT | β' | n.d. | n.d. | n.d. | n.d. |
| GVKLHYT | β' | n.d. | n.d. | n.d. | n.d. |
| GVKLHYE | β' | 2.3 ± 0.23 | 2424.7 ± 178.3 | 12.7 ± 0.3 | 0.331 |
| KEVYLHG | β' | n.d. | n.d. | n.d. | n.d. |
| GVKLHYT | β'-Tyr33Ala | n.d. | n.d. | n.d. | n.d. |
| GVKLHYE | β'-Tyr33Ala | n.d. | n.d. | n.d. | n.d. |
| GVKLHYT (amidated) | α | n.d. | n.d. | n.d. | n.d. |
| GVKLHYE (amidated) | α | 2.3 ± 0.7 | 706.2 ± 56.3 | 8.8 ± 0.2 | 0.01 |
| GVKLHYT | α-Tyr139Ala | n.d. | n.d. | n.d. | n.d. |
| GVKLHYE | α-Tyr139Ala | $3.8 \pm 0.36$ | $160.8 \pm 15.8$ | $4.4 \pm 0.15$ | 0.154 |
| KEVYLHG | α-Tyr139Ala | n.d. | n.d. | n.d. | n.d. |
| GVKLHYT | α-Arg13Ala | $8.5 \pm 2.0$ | $147.7 \pm 7.0$ | $3.8 \pm 0.07$ | 0.033 |
| GVKLHYE | α-Arg13Ala | $3.0 \pm 0.2$ | $132.5 \pm 4.5$ | $2.2 \pm 0.08$ | 0.316 |
| GVKLHYT | α-Arg300Ala | $3.6 \pm 0.4$ | $360.1 \pm 9.4$ | $8.1 \pm 0.07$ | 0.139 |
| GVKLHYE | α-Arg300Ala | $4.2 \pm 0.4$ | $136.6 \pm 2.6$ | $4.3 \pm 0.03$ | 0.664 |
| GVKLHYT | α-Lys15Ala | n.d. | n.d. | n.d. | n.d. |
| GVKLHYE | α-Lys15Ala | n.d. | n.d. | n.d. | n.d. |
| CKFDEDDSEPVLKGVKLHYT | $\alpha$ | $1.7 \pm 0.083$ | $99.3 \pm 1.5$ | $2.67 \pm 0.01$ | 0.854 |
| CKFDEDDSEPVLKGVKLHYE | $\alpha$ | $0.1 \pm 0.005$ | $59.9 \pm 1.3$ | $0.037 \pm .002$ | 0.415 |
| CKFDEDDSEPVLKGVKLHYT<sup>2</sup> | $\alpha$ | $1.1 \pm 0.1$ | $145.5 \pm 4.4$ | $2.9 \pm 0.03$ | 0.815 |
| CKFDEDDSEPVLKGVKLHYE<sup>2</sup> | $\alpha$ | $0.1 \pm 0.002$ | $60.8 \pm 1.3$ | $0.03 \pm 0.002$ | 0.423 |

<sup>1</sup>Lys1269 and His1271 are highlighted in cyan, and Glu1273 in red
<sup>2</sup>These assays were performed at pH 7.0 as opposed to pH 6.5 for other assays.

### The clientized S tail has broad selectivity for αWD40 and β’WD40

Prior work revealed that the wild-type S heptapeptide does not bind to β’WD40^34^, even though there is complete conservation of the basic cluster in α and β’WD40^43,44^. We therefore asked whether clientization of the S heptapeptide enables binding to β’WD40, and indeed, clientized S heptapeptide bound to purified β’WD40 with a K_D_ of 2.30 ± 0.23 μM **(Table 1, Figure 3A, B)**. This suggests that the x_CT_ residue, which is outside the dibasic motif, is a critical determinant of S-coatomer selectivity.

**Figure 3.**
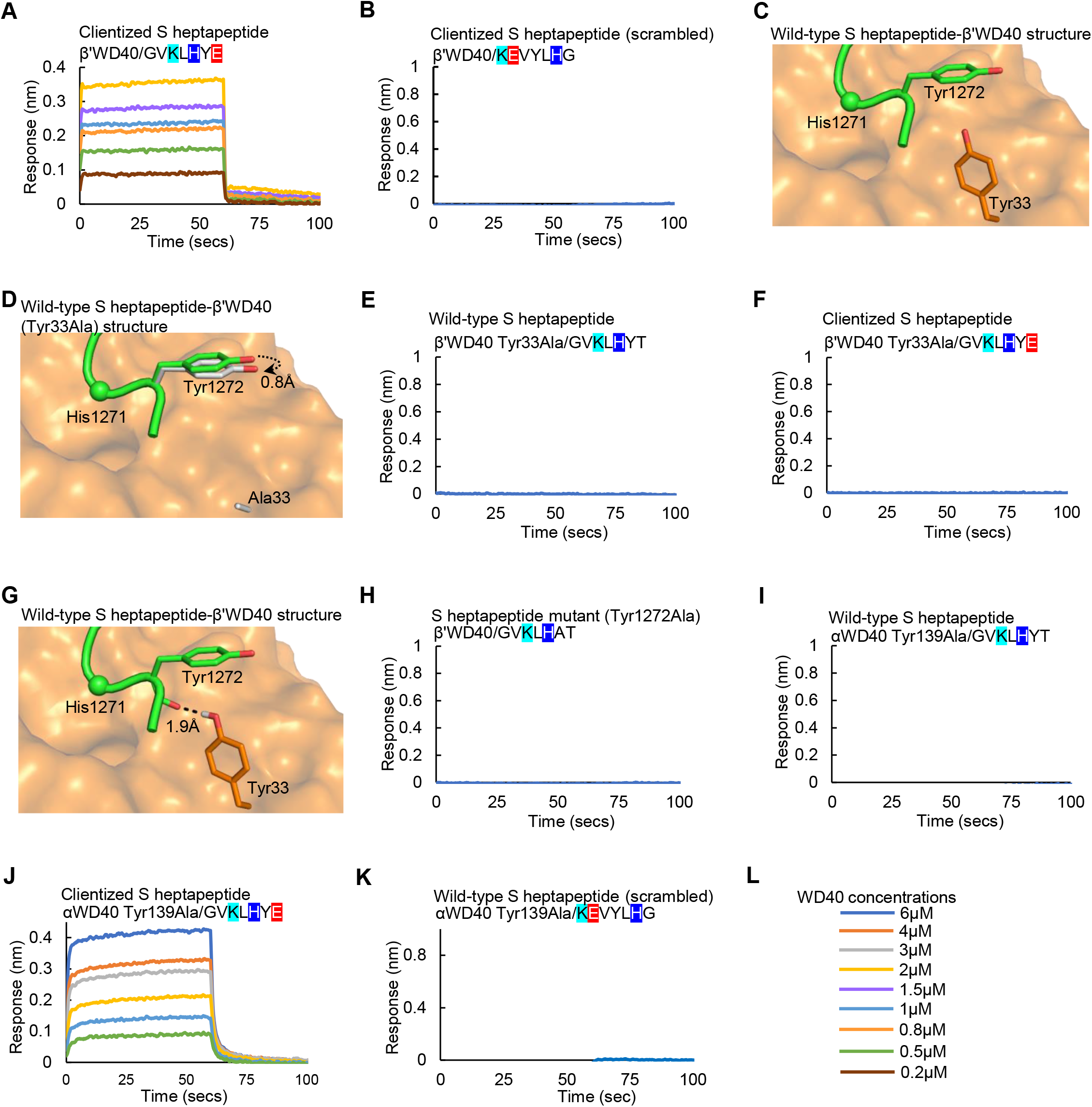
The clientized S tail has broad selectivity for αWD40 and β’WD40 domains. **(A, B)** BLI assays show direct binding of clientized S heptapeptide to β’WD40 **(A)** and no binding for a clientized S heptapeptide with scrambled sequence **(B). (C, D)** Co-crystal structures show the β’WD40 Tyr33 side chain pushing the clientized S heptapeptide Tyr1272 side chain away from its surface **(C)** and substitution of the β’WD40 Tyr33 side chain for Ala causing minimal movement of the S tail Tyr1272 side chain towards its surface **(D). (E, F)** In BLI assays, β’WD40 Tyr33Ala mutant does not bind to either wild-type **(E)** or clientized **(F)** S heptapeptide. **(G)** Co-crystal structure of β’WD40 with clientized S heptapeptide shows Tyr33 providing a main chain interaction that is likely disrupted in the Tyr33Ala mutant. **(H-K)** In BLI assays, the Tyr1272Ala S heptapeptide fails to bind to β’WD40 **(H)** and the wild-type S heptapeptide fails to bind to Tyr139Ala αWD40 **(I)**. Clientized S heptapeptide binds to αWD40 Tyr139Ala **(J)** but not when the clientized S heptapeptide sequence is scrambled **(K)**. Color key indicates WD40 concentrations for BLI assays.

Next, we asked why the wild-type S heptapeptide does not bind to wild-type β’WD40. Our co-crystal structures showed that the Tyr33 side chain of β’WD40 pushes away the Tyr1272 side chain of the S heptapeptide **(Figure 3C)**. Hence, we hypothesized that replacement of the bulky Tyr33 side chain with a smaller residue would allow Tyr1272 to move closer to the β’WD40 surface, strengthening binding between S and β’WD40. We generated a β’WD40 Tyr33Ala mutant and determined its co-crystal structure with the clientized S heptapeptide at a resolution of 1.8 Å **(Extended Data Table S2)**. Although the S Tyr1272 side chain was closer to the mutant β’WD40 surface by 0.8 Å **(Figure 3D; Extended Data Figure S2F**), the Tyr1272 side chain was still > 4 Å away from the β’WD40 surface. A BLI assay revealed this was insufficient to enhance S-β’WD40 affinity **(Figure 3E)**. Moreover, this β’WD40 Tyr33Ala mutation abolished binding to the clientized S heptapeptide **(Figure 3F)**, suggesting a previously unrecognized role for β’WD40 Tyr33 in S heptapeptide binding.

It seemed likely that the Tyr33Ala mutation disrupted a hydrogen bond between the Tyr33 hydroxyl group on β’WD40 and the main chain carbonyl O atom of Tyr1272 on the S heptapeptide **(Figure 3G)**. To test the role of steric hindrance at this Tyr1272-Tyr33 interaction site without disrupting such stabilizing interactions, we mutated Tyr1272 on the heptapeptide to alanine (1267Gly-Val-<u>Lys</u>-Leu-<u>His</u>-Ala-Thr1273), but there was no enhancement to β’WD40 binding **(Figure 3H)**. We therefore asked if affinity for the wild-type S heptapeptide requires not only elimination of steric hindrance at the 33^rd^ residue of β’WD40, but also the introduction of stabilizing interactions. Such potentially stabilizing interactions are offered by αWD40 His31, which is juxtaposed with β’WD40 Tyr33. Therefore, we utilized an αWD40 mutant that is a structural and functional chimera of αWD40-β’WD40. The Tyr139 residue in this αWD40 mutant is replaced with an alanine. This generates functional equivalence at this site as neither Ala139 in this αWD40 mutant nor the juxtaposed β’WD40 Phe142 provides hydrogen bonding to S^34^. Furthermore, this αWD40 mutant retains the native His31 residue, which is juxtaposed with β’WD40 Tyr33, and provides hydrogen bonding to S^34^. Finally, the binding site of this mutant has a similar architecture to wild-type β’WD40^34^. In BLI assays, we found specific binding of this αWD40 mutant to the clientized S heptapeptide with a K_D_ of 3.80 ± 0.36 μM, unlike the non-binder wild-type S heptapeptide **(Figure 3I-K)**. Together, these assays show that S-WD40 binding involves a combination of bonding and steric interactions with the juxtaposed αWD40 His31 and β’WD40 Tyr33 residues, and that loss of critical interactions at upstream sites can be overcome by clientizing the S x_CT_ residue. Hence, the S protein C-terminus is a critical modulator of coatomer interactions.

### Clientization alters the S tail conformation and strengthens binding to WD40 domains

Because the clientizing mutation Thr1273Glu generates a side chain and main chain anionic cluster in a highly charged S tail **(Extended Data Figure S1A)**, we sought to determine whether clientization induces a distinct conformation. We first assigned the NMR spectra of a ^13^C/^15^N-labeled wild-type S tail **(Figure 4A)**. Comparison of ^13^Cα and ^13^CO chemical shifts with corresponding calculated random coil values indicated that Glu1258-Ser1261 and Val1268-His1271 had a weak propensity to form extended β-strand-like structures **(Figure 4B)**. However, wild-type and clientized C-terminal residues showed large differences in chemical shift perturbations (CSPs), suggestive of conformational differences in this region **(Extended Data Figure S3)**.

**Figure 4.**
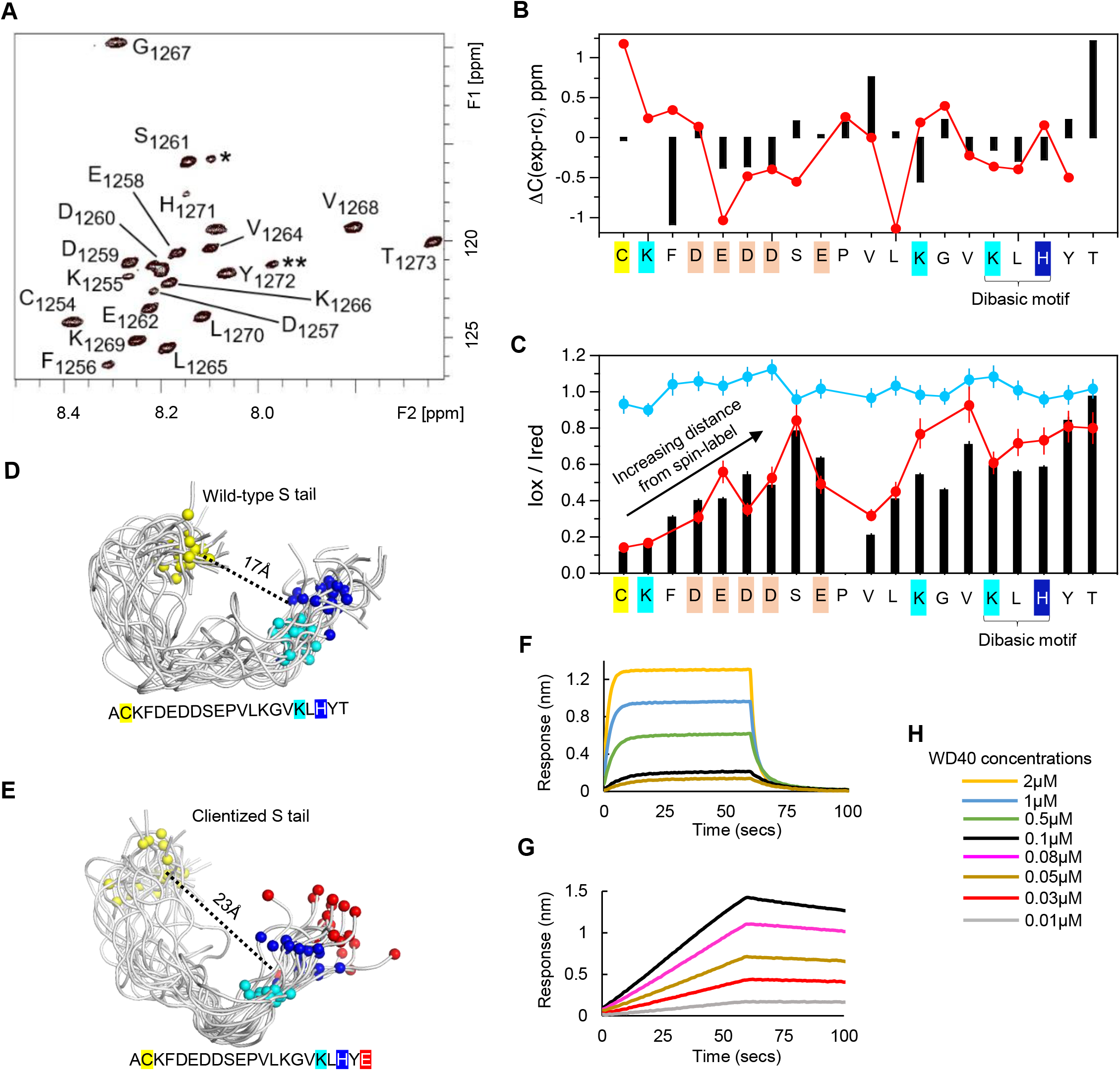
Clientization alters the S tail conformation and strengthens binding to WD40 domains. **(A)** Two-dimensional ^1^H-^15^N HSQC spectrum of ^15^N-isotope labelled wild-type S tail 21-mer peptide at 25^°^C with amide assignments. Peaks marked * and ** are likely due to Ser1261 and Glu1262, respectively, in a small population of peptide with cis-Pro1263. F2 = ^1^H; F1 = ^15^N. **(B)** Plot of ^13^C chemical shift differences (experiment-calculated random coil) in the ^13^C/^15^N-SARS-CoV-2 S tail peptide for Cα (black) and CO (red) resonances. Conformational preferences for extended strand-like structures is observed in residues Glu1258-Ser1261 and the dibasic motif-containing Val1268-His1271. **(C)** PRE plots of I_ox_/I_red_ versus residue for wild-type ^15^N-SARS-CoV-2 S tail 21-mer peptide (black), Thr1273Glu mutant (red), and control for intermolecular effects (light blue). S tail residues highlighted as follows: Cys (yellow), Lys (cyan), His (dark blue), acidic residues (tan). **(D, E)** NMR ensembles of wild-type **(D)** and clientized **(E)** S tails, highlighting Cα atoms for Cys1254 (yellow), Lys1269 (cyan), His1271 (dark blue), and Glu1273 (red). The clientized tail has a larger distance between Cys1254 and His1271. **(F, G)** BLI assays showing αWD40 interactions of wild-type **(F)** and clientized **(G)** S tails, the latter being substantially stronger but slower. **(H)** Concentration of WD40 domains used in BLI assays.

To directly evaluate conformational changes induced by the clientizing mutation, we used paramagnetic relaxation enhancement (PRE)^45,46^. The spin label MTSL (S-(1-oxyl-2,2,5,5-tetramethyl-2,5-dihydro-1H-pyrrol-3-yl)methyl methanesulfonothioate) was conjugated to a single cysteine at position 1254 near the N-terminus of the S tail. Amide peak intensities in the 2D ^1^H-^15^N heteronuclear single quantum coherence (HSQC) spectrum of this ^15^N-isotope labelled and MTSL-conjugated S tail were measured in the paramagnetic (I_ox_) and reduced (I_red_) state using ascorbate as a diamagnetic control **(Figure 4C)**. The paramagnetic broadening of backbone amide protons obtained from I_ox_/I_red_ intensity ratios provides long-range (< 25 Å) distance constraints between the MTSL spin label and amide protons^45^. Therefore, PRE can only be detected up to ∼ 8 residues from the spin label in a random coil. As expected, PRE effects progressively attenuated with distance from the N-terminal spin-label up to Ser1261, but lower I_ox_/I_red_ values indicated increased PRE between residues Glu1262 and His1271. This suggests a hinge-like structure centered around Glu1262 and Pro1263 and consequent masking of the C-terminal dibasic motif by the N-terminal residues in the wild-type S tail. In contrast, the clientized S tail showed significantly higher I_ox_/I_red_ values for Val1264, Lys1266, Val1268, Leu1270, and His1271 (**Figure 4C**), indicating that the clientizing mutation weakens transient contacts between the N- and C-terminal halves.

We used these PRE data as restraints to generate NMR ensembles of wild-type and clientized S tails **(Figure 4D, E)**. This revealed a ∼ 6 Å larger separation between the N-terminal Cys1254 residue and the dibasic motif, as well as a more solvent-accessible extended motif in the clientized S tail relative to wild type. We thus hypothesized that binding of the clientized S tail to WD40 domains will be faster and stronger than the more compact wild-type tail. To test this hypothesis, we determined the relative binding of full-length, extra-membrane S tails to αWD40 and β’WD40 domains. Wild-type S tail CSPs were substantially smaller for β’WD40 than αWD40 **(Extended Data Figure S4)**, indicating weaker binding to β’WD40, but clientized S tail CSPs revealed enhanced binding to β’WD40 **(Extended Data Figure S5)**. These data validated our findings from shorter S tail hepta-peptides. Furthermore, a BLI analysis showed 11-fold tighter binding of the full-length clientized S tail to αWD40, relative to wild type **(Figure 4F, G, Table 1)**. Interestingly, the αWD40 binding rate was 2.4-fold slower in the clientized S tail. Because superposition of WD40-S tail co-crystal structures on S tail NMR ensembles show modest differences in clashes between WD40 and NMR-resolved tail residues for both wild-type and clientized S tails **(Extended Data Table S3)**, we infer that slower binding of the clientized S tail is likely due to a modulatory role of the Glu1273 side chain rather than tail conformation.

Our NMR experiments identified S tail residues well upstream the crystallographic footprint that demonstrate CSPs upon binding to αWD40 (**Figure 4C; Extended Data Figure S4**). These CSPs could be due to direct contacts between the S tail and WD40 domain, reflecting a larger binding footprint for the full-length S tail, or indirect effects such as a loss of long-range intra-tail contacts upon binding to αWD40. Interestingly, the upstream residues Ser1261-Glu1262, which modulate coatomer interactions^35^, are in a turn that likely modulates N- and C-terminal interactions in the S tail **(Extended Data Figure S6)**. Nevertheless, these various data show that clientization enhances the binding of S proteins to WD40 domains.

### Enhanced coatomer affinity reduces S-directed membrane fusion

Having established that clientization of the S protein enhances coatomer binding and localization in early secretory compartments, we asked whether it might also modify export to the host cell’s PM as well as fusogenicity. To address these questions, we performed a dual split protein (DSP) assay in cell culture^47^ **(Figure 5A)**. In this assay, the S protein is on the PM of “effector” cells, which express one half of a split luciferase. The receptor ACE2 is on “target” cells, which express the other half of the split luciferase. S-ACE2 interaction and subsequent fusion are followed by the reconstitution of the luciferase, whose activity serves as a read-out for S protein fusogenicity. In contrast to wild-type S protein, which yielded a robust luciferase signal, clientized S protein containing either a Thr1273Glu or Thr1273Asp mutation showed no significant enhancement of luciferase activity **(Figure 5B)**. Moreover, western blot analysis of effector cell lysates revealed an intact, unproteolyzed population of clientized S protein, revealing a lack of trafficking from early secretory compartments **(Figure 5C)**. This further suggests that M, E, and N structural proteins are unable to liberate clientized S from coatomer for export to the PM.

**Figure 5.**
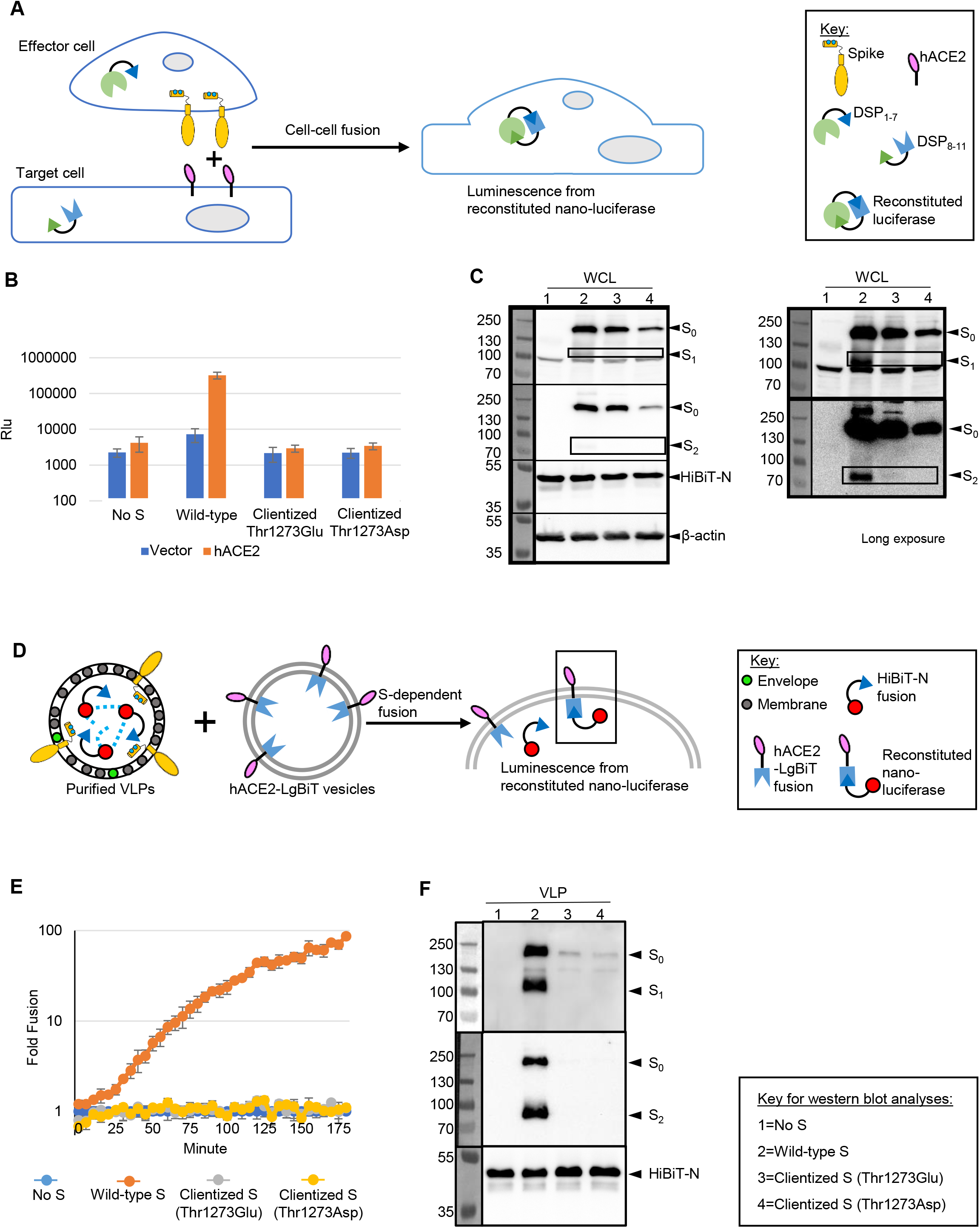
Clientization inhibits S protein fusogenicity and incorporation in VLPs. **(A)** Schematic of S protein fusogenicity assay. If fusion-competent S proteins traffic to the surface of S-expressing HEK293T (effector) cells, they engage with HEK293T (target) cells expressing the human receptor for SARS-CoV-2 (hACE2) and activate cell-cell fusion. This results in multi-nucleate syncytia, allowing DSP reconstitution of nano-luciferase from DSP_1-7_ and DSP_8-11_. Reconstitution does not occur if S is fusion incompetent or not trafficked to the PM. **(B)** Nano-luciferase luminescence for wild-type, and clientized S in protein fusogenicity assays. Clientized S shows 10-fold lower nano-luciferase luminescence than wild-type S. Data represent mean and standard error. **(C)** Western blots of whole cell lysates (WCLs) from effector cells used for protein fusogenicity assays show intact S (S_0_) from early secretory compartments and cleaved S fragments (S_1_ and S_2_) from PM when wild-type S is expressed. In contrast, no proteolyzed fragments are observed when clientized S is expressed. Right panel: Longer exposure western blot analysis. **(D)** Schematic of SARS-CoV-2 VLP assembly assay. S, M, E, and HiBiT-N plasmids co-transfected in HEK293T cells generate VLPs, which are purified from the extracellular medium. VLPs displaying functional S proteins will engage vesicles displaying hACE2 fused to a cytosolic LgBiT fragment, resulting in VLP-vesicle fusion, nano-luciferase reconstitution, and a luminescence signal. **(E)** Luminescence signals from VLP assembly assay. Only wild-type S results in fusogenic activity and enhanced luminescence. Data represent mean and standard error. **(F)** Western blots of purified VLPs show the abundant presence of wild-type S and the absence of clientized S.

Next, we asked whether intracellular retention of clientized S increases its abundance at SARS-CoV-2 budding sites, conceivably promoting progeny infectivity, or whether strong coatomer association inhibits incorporation of S into assembling virions. In the latter scenario, CoV particles will have a lower abundance of S proteins and thus diminished cell entry potential. We employed a SARS-CoV-2 virus-like particle (VLP) assembly system that utilized luminescence as an assay for S-dependent entry of VLPs^47,48^ **(Figure 5D)**. VLPs secreted from cells containing wild-type S protein generated robust luminescence and contained abundant furin-cleaved S **(Figure 5E, F)**. In sharp contrast, VLPs from cells expressing clientized S failed to generate luminescence and contained few S molecules **(Figure 5E, F)**. Thus, strong association between clientized S and coatomer interferes with incorporation of S into newly assembling VLPs and compromises VLP-directed membrane fusion.

### The retrograde pathway traffics S protein to the virion assembly site

Finally, to determine whether the anterograde or retrograde pathway is the primary conduit for S protein during SARS-CoV-2 assembly, we utilized a Lys1269Ala/His1271Ala double mutant of S dibasic motif, which is unable to interact with coatomer^35,36^. This retrograde trafficking-defective S mutant instead undergoes selective anterograde trafficking. VLPs secreted from cells containing this S mutant resulted in ∼ 75% less membrane fusion than VLPs containing wild-type S **(Figure 6A)**. Moreover, these retrograde trafficking-defective S molecules were less efficiently incorporated into VLPs **(Figure 6B, C)**. Interestingly, retrograde trafficking-defective S protein caused 2.3-fold higher fusogenicity of cells in culture than wild-type S protein **(Figure 6D)**, consistent with its facile escape from the coatomer and consequent population of the PM.

**Figure 6.**
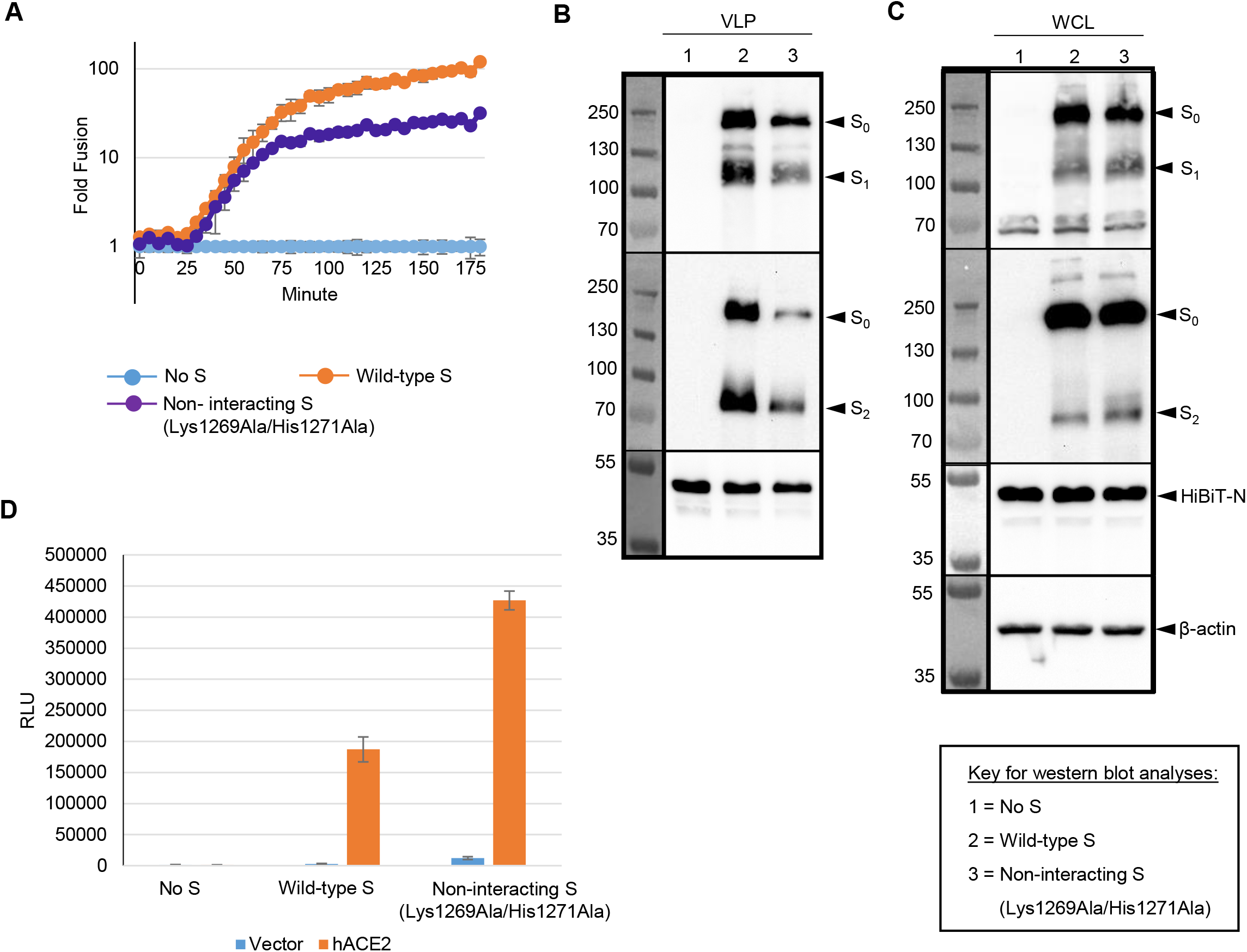
Coatomer binding facilitates incorporation of S into VLPs and inhibits cell-cell fusogenicity. **(A)** Luminescence signals from VLP assembly assay. An S protein double mutant (Lys1269Ala/His1271Ala) that does not interact with coatomer shows a substantially smaller increase in luminescence than wild-type indicating poor incorporation into VLPs. Data represent mean and standard error. **(B)** Western blots of purified VLPs confirm poor incorporation of the non-interacting S protein. **(C)** Western blots of WCLs show no expression differences between the wild-type and non-interacting S protein. **(D)** Nano-luciferase luminescence for non-interacting S protein is 2.3-fold higher than wild-type. Data represent mean and standard error.

## DISCUSSION

In this study, we used structure-function analyses to determine the molecular function of the non-acidic C-terminal residue of the SARS-CoV-2 S protein, revealing the importance of this lack of mimicry of coatomer interacting residues. Our work revealed the atomic basis of the S-coatomer interaction as well as fundamental details of S protein trafficking and progeny assembly.

In particular, our investigation showed that enhancing mimicry of the client protein extended motif by introducing an acidic residue at Thr1273 substantially increases coatomer binding, thereby localizing mutant S proteins in coatomer-enriched early secretory compartments. Furthermore, this manipulation provided a means to show that the balance between coatomer-dependent retention and release of S protein is critical for SARS-CoV-2 propagation. If the S-coatomer interaction is enhanced, virion assembly and cell-cell fusion are greatly reduced. Alternatively, if the S-coatomer interaction is abolished, virion fusogenicity is reduced while cell-cell fusion is increased due to greater anterograde S transport. Hence, an intermediate S-coatomer interaction affinity provides a trade-off between virion assembly and cell-cell fusion, allowing both virus transmission within and between host animals. As such, a single residue at the C-terminus of the S tail is a critical determinant of SARS-CoV-2 propagation.

We also elucidated the structural and biophysical basis of the balance between binding and dissociation of the S protein and coatomer. We showed that clientization of the S protein by mutation of Thr1273 to glutamate generates an electrostatic lock between the acidic side chain of glutamate and a cluster of two basic residues conserved in α and β’WD40. The main chain carboxylate of Thr1273 and of Glu1273 interacts with Lys15, the third residue in this basic cluster. Site-directed mutagenesis of this basic cluster showed that an arginine at position 300 in αWD40 determines this enhanced affinity. In contrast, Arg13 and Lys15 residues in αWD40 provide the basis for basal binding to both wild-type and clientized spikes. Hence, the WD40 basic cluster has evolved distinct and fine-tuned interaction sites that modulate binding affinities for client proteins and the CoV S protein. Our analysis thus reveals key atomic-level selectivity principles in coatomer-client interactions. In particular, the C-terminal acidic residue in client proteins modulates coatomer subunit binding affinity, and hence selectivity. We infer that this subunit selectivity is not absolute but instead involves relative modulation of interaction affinity for one coatomer subunit versus the other. Furthermore, our data suggest that simultaneous, high avidity engagement of α and β’-COPI subunits is likely driven by the C-terminal residue in endogenous oligomeric clients such as the ER-resident enzyme UGT^49^, which is critical for xenobiotic processing and displays a dibasic motif in its cytosolic tail. Because more than 100 enzymes, signaling proteins, and growth factors in the eukaryotic proteome display a dibasic motif^34^, the structural principles of coatomer-client affinity modulation that we have elucidated have broad implications for secretory homeostasis.

Our analyses of SARS-CoV-2 VLP assembly revealed that incorporation of retrogradely trafficked S protein is favored over newly synthesized protein delivered by anterograde trafficking from the ER. Golgi-associated S protein may be identified for retrograde trafficking by its association with Golgi-resident CoV M proteins^1,2^, consistent with its dependence on cis-Golgi-located M protein for incorporation into virions^2,8^. However, because the CoV M protein lacks a dibasic retrieval motif, our data suggest that the S protein acts as a chaperone to deliver M to ERGIC. Once delivered, M protein may direct S into budding virions. Our investigation also raises intriguing questions about the balance between anterograde and retrograde trafficking of S at various stages of infection. Presumably, newly synthesized S is at low concentration during the early stages of infection and therefore efficiently retrieved by the coatomer for virion assembly. However, higher concentrations of S in later stages are likely to saturate the coatomer, resulting in export of S to the PM for cell-cell-fusion. This raises the interesting hypothesis that a time delay between infection and cell-cell fusion ensures progeny are assembled prior to transmission. Both this time delay and the evolutionary specialization of M and S proteins requires further investigation.

Although retrograde and anterograde trafficking of the S protein are critical for SARS-CoV-2 propagation, the mechanism underlying trafficking bifurcation at the cis-Golgi is not well understood. Structural insights from our NMR studies advance the understanding of this intriguing mechanism. The S tail displays extensive conformational changes that modify exposure of the extended motif, suggesting a mechanism wherein access to the extended motif modulates coatomer binding. S molecules in the cis-Golgi with an inaccessible extended motif will be incompetent for coatomer binding and continue anterograde export to the PM. In contrast, an accessible extended motif will permit S retrieval to the virion assembly site in ERGIC. Entire localization of clientized S protein in early secretory compartments suggests that the Thr1273Glu mutation alters this conformational equilibrium, favoring states with an accessible extended motif. This would enable efficient retrieval of S protein by the coatomer and reduce the probability of anterograde export to the PM. Such an equilibrium in S tail states might be modulated by factors such as oligomerization, palmitoylation, proximity to the Golgi membrane, association with cytosolic proteins, and mutations outside the extended motif^35,50,51^. For example, mutagenesis of the SARS-CoV-2 S tail has identified residues upstream of the extended motif (Ser1261-Glu1262) that are critical for coatomer interactions^35^. Our structural analysis maps these two residues to a turn in the S tail, suggesting that the Ser1261Ala/Glu1262Ala double mutation modifies the conformational accessibility of the extended motif and reduces coatomer binding. On a broader level, our analysis reveals that indirect, long-range conformational dynamics can determine S-coatomer binding and the balance of anterograde-retrograde trafficking.

Bifurcation of S trafficking has implications for vaccinology. Our data suggest that the C-terminal residue of S protein encoded by mRNA or adenoviral constructs^4^ could modulate S export and its antigenic display on the cell surface. Such C-terminus optimization will be essential for S export and a robust immune response to genetic vaccines against the wide diversity of C-terminal residues in S proteins encoded by CoVs of human concern, including the lethal MERS-CoV and human CoV OC43 associated with common cold-like symptoms^34,52^. Furthermore, our research raises important questions about the modulation of endogenous protein trafficking upon transient expression of S-based genetic vaccines, as disruption of coatomer function has serious clinical outcomes. For instance, disruption of client retrograde retrieval due to mutations in COPA, the human ortholog of α-COPI, underlies the autoimmune COPA syndrome, which causes coughing, shortness of breath, lung inflammation and scarring, pulmonary hemorrhaging, and systemic autoimmune reactions such as arthritis^53^. This emphasizes an urgent need to investigate secretory disruptions due to coatomer hijacking by SARS-CoV-2 or by transient expression of S-based genetic vaccines.

In summary, we have shown that the bipartite organization of the S protein extended motif balances two opposing forces: coatomer hijacking to traffic S to the ERGIC and S-coatomer dissociation at this assembly site. Together with histone mimicry by SARS-CoV-2 to modify the host epigenome and anti-viral response^54^, our investigation establishes molecular mimicry as a major mechanism employed by SARS-CoV-2 to achieve infection and progeny assembly.

## DATA AVAILABILITY

All crystal structures have been deposited in the Protein Data Bank and are listed in Extended Data Table T2. All plasmids will be deposited in Addgene.

## METHODS

### Immunofluorescence microscopy

For determination of the subcellular localization of either wild-type S protein or the Thr1273Glu mutant, the two DNA constructs were transfected into HeLa cells using jetOPTIMUS transfection reagent. These cells were on sterile glass coverslips. Fixation was performed the next day with 4% formaldehyde (Sigma-Aldrich) for 10 min. Then, permeabilization and and blocking were performed with PBS (with 0.4% (v/v) Triton X-100%) and 2% immunoglobulin G-free BSA (Jackson ImmunoResearch) for 1 hour. The anti-spike 1A9 mouse monoclonal antibody (Cat #GTX632604) from GeneTex (Irvine, CA, USA) in PBS (with 0.1% Triton X-100% and 0.5% BSA) was used to probe the cells. These cells were then mounted in ProLong® Glass antifade mounting medium (Life Technologies) following treatment with fluorophore-conjugated secondary antibody and washing. An LSM880 confocal microscope (Carl Zeiss Inc., Peabody, MA) was used to acquire images, which were then analyzed by ImageJ software (Fiji).

### Preparation of mouse liver cytosol (MLC)

Livers extracted from freshly killed C57Blk6 mice were used for MLC. Livers (10 grams) were chopped into small pieces on a petri plate on ice and homogenized with 20ml homogenization buffer A (25 mM HEPES-KOH pH 7.4, 125 mM potassium acetate, 2.5 mM magnesium acetate, 1 mM DTT) using a Dounce glass homogenizer (50 strokes). The homogenized extract was first spun at 800 xg for 10 mins at 4 °C to remove the nuclei, then again at 10,000 xg for 10 mins at 4°C to remove the mitochondria. The resulting supernatant was then subject to ultracentrifugation at 100,000 xg at 4°C to separate membranes from cytosol. The clarified cytosolic fraction was then carefully removed and stored in 1 ml aliquots at -80°C.

### GST pull-down assays

For GST pull-down assay, a tube of the 1 ml mouse liver cytosol was thawed on ice, centrifuged at 20,000 xg at 4°C to remove any precipitated material, and the clarified cytosol was then diluted to approximately 20 mg/ml with cold homogenization buffer B (homogenization buffer A containing TX-100 at a final concentration of 0.1%). For binding assays, 100 μg of each purified fusion protein was first immobilized on 50 μl of packed glutathione-agarose beads at room temperature for 1 hour, the beads were washed once with cold homogenization buffer B, then 300 μl of the diluted cytosol was added and the beads tumbled for 2 h at 4°C. The beads were washed 3 times with cold homogenization buffer B, and the pellets resuspended in sample buffer and heated at 100°C for 10 mins before SDS-PAGE.

### Western blot and antibodies

In Figures 5 and 6, samples in SDS solubilizer [0.0625 M Tris·HCl (pH 6.8), 10% glycerol, 0.01% bromophenol blue, 2% (wt/vol) SDS, +/- 2% 2-mercaptoethanol] were heated at 95°C for 5 min, electrophoresed through 8% or 10% (wt/vol) polyacrylamide-SDS gels, transferred to nitrocellulose membranes (Bio-Rad), and incubated with rabbit polyclonal anti-SARS-CoV-2-S1 (SinoBiological, cat: 40591-T62), mouse monoclonal anti-SARS-S2 (ThermoFisher, cat: MA5-35946, conjugated to HRP), goat anti-human IgG (sc-2453, Santa Cruz Biotechnologies), rabbit monoclonal anti-hACE2 (Invitrogen, cat: MA5-32307), or purified LgBiT-substrate cocktail (Promega). After incubation with appropriate HRP-tagged secondary antibodies and chemiluminescent substrate (Thermo Fisher), the blots were imaged and processed with a FlourChem E (Protein Simple). In Figure 1, the anti-spike 1A9 mouse monoclonal antibody (Cat #GTX632604) from GeneTex (Irvine, CA, USA) was used in the western blot analysis.

### Coatomer WD40 expression and purification

Recombinant expression and purification of *S. pombe* αWD40 and *S. cerevisiae* β’WD40 domains (both wild type and mutants) were carried out as described ^34,55^. Briefly, αWD40 with a C-terminal strep-tag was expressed and purified from Expi293 cells using polyethyleneimine (PEI). Clarified cell lysate was subjected to affinity chromatography followed by size-exclusion chromatography (SEC) on a preparative grade Superdex-75 column.

β’WD40 domain with an N-terminal strep-Hisx6-SUMO tag was expressed in BL21(DE3)pLysS cells by IPTG induction at 18°C overnight. After affinity chromatography, β’WD40 was further subjected to ULP1 protease digestion for removal of the SUMO tag. Further purification was performed Ni-NTA chromatography and SEC. Plasmids for all WD40 mutants (αWD40 Arg13Ala, Lys15Ala, Arg300Ala, and β’WD40 Tyr33Ala) were generated using the Q5® Site-Directed Mutagenesis Kit and were purified as described above.

### Biolayer interferometry (BLI)

BLI assays with biotinylated S peptides from Biomatik and purified coatomer WD40 domains were carried out as described ^34^. Briefly, WD40 was provided as the analyte with a buffer composition of 20 mM Tris-HCl (pH 7.5) or 50 mM MES-NaOH (pH 6.5), 150 mM NaCl, 5 mM DTT, 10% glycerol, 0.2 mg/ml bovine serum albumin (BSA), and 0.002% Tween 20. S peptides were immobilized on streptavidin (SA) biosensors (FortéBio). Measurements were performed on the Octet RED96 system (FortéBio). Data acquisition and data analysis were performed using Data Acquisition 11.1 suite and FortéBio Data Analysis 11.1 software suite respectively. All experiments were performed at 25°C and 1000 rpm shake speed. Appropriate sensor and loading controls were applied.

### X-ray crystallography

Crystallization was performed as described ^34^. Briefly, purified mutant αWD40 domains were concentrated to 2mg/ml. Crystal trays were set-up using the hanging drop vapor diffusion method with αWD40 domains mixed with reservoir buffer in 1:1 v/v ratio. Crystals grew at room temperature within 24 hours. Crystals of αWD40 Arg13Ala grew in 0.14M sodium citrate tribasic dehydrate, 16% PEG3350. Crystals of αWD40 Lys15Ala grew in 0.14M sodium citrate tribasic dihydrate, 18% PEG3350. Crystals of αWD40 Arg300Ala grew in 0.2M potassium sodium tartrate tetrahydrate, 18%PEG3350.

The purified β’WD40 domain was concentrated to 20-25 mg/ml. Co-crystals of β’WD40 domain with wild-type and clientizing S hepta-peptides were set-up with a protein:peptide molar ratio of 1:2. Co-crystals grew in 2-4 days at room temperature. Wild-type β’WD40 with wild-type S hepta-peptide grew in 0.1M MES pH 6.2, 19% PEG 4000. Wild-type β’COPI WD40 with clientizing S hepta-peptide grew in 0.1M MES pH 6.0, 15% PEG20000. Co-crystals of β’WD40-Tyr33Ala mutant with clientizing S hepta-peptide grew in 0.1M MES pH 6.2, 17% PEG20000.

X-ray diffraction data collection was carried out at the National Synchrotron Light Source II (NSLS II) beamline 17-ID-1 AMX at the Brookhaven National Laboratory. Diffraction data was indexed, integrated, and scaled in XDS ^56^ as part of the beamline data acquisition and processing pipeline. Data processing statistics are outlined in **Extended Data Table T2**. Molecular replacement was performed in Phenix using PDB ID’s 7S22 for αWD40 mutants and 4J79 for β’WD40 in complex with S hepta-peptides ^34,44,57^. Refinement and model building was performed using the Phenix.refine module and Coot. All figures were generated using Pymol and Coot.

### Preparation of isotope-labeled S tail peptides

A gene fragment corresponding to the wild-type SARS-CoV-2 S C-terminal tail was cloned into an eXact tag pH720 vector system ^58^. Isotope-labeled (^15^N, ^13^C/^15^N) peptide was prepared by transforming the construct into *E. coli* BL21(DE3) cells, followed by growth in M9 minimal media with either ^15^N-ammonium chloride or ^13^C-glucose/^15^N-ammonium chloride at 37°C until an OD600∼0.6-0.9 was reached. Protein expression was induced with 1 mM IPTG for 18 hours at 25°C. Cells were centrifuged, re-suspended, and lysed using sonication. The cleared lysate was purified on an immobilized subtilisin column (Potomac Affinity Proteins), using a method similar to that described previously ^59^. Fractions containing at least 95% pure peptide, as determined by MALDI, were pooled and concentrated for further analysis. The Q5 site-directed mutagenesis kit (New England Biolabs) was used for preparation of mutants.

### NMR spectroscopy

NMR spectra were acquired on a Bruker Avance III 600 MHz spectrometer fitted with a Z-gradient ^1^H/^13^C/^15^N-cryoprobe. Backbone resonance assignments for the 21-residue SARS-CoV-2 S C-terminal tail peptide were made using heteronuclear triple resonance experiments as follows: HNCACB, CBCA(CO)NH, HNCO, HN(CA)CO, and (H)N(CA)NNH. Sample conditions were 300 μM peptide, 100 mM KPi, 1 mM DTT, pH 7.0, 25°C. Secondary shifts were calculated from experimental and sequence corrected random coil chemical shifts ^60^. Spectra were processed with NMRPipe ^61^ and analyzed with Sparky ^62^.

Chemical shift mapping experiments were performed by comparing two dimensional ^1^H-^15^N HSQC spectra of ^15^N-labeled SARS-CoV-2 S tail in its unbound and WD40-bound states. Bound states were prepared by adding unlabeled αWD40 or β’WD40 to ^15^N-labeled peptide with a WD40/peptide molar ratio of ∼1.2:1. Chemical shift perturbations were calculated from the equation, Δδ_total_ = [(*W*_H_Δδ_H_)^2^+(*W*_N_Δδ_N_)^2^]^1/2^, where WH =1 and WN = 0.2 ^59^. To allow for more direct comparison with the BLI results, sample conditions for shift mapping were 20 mM Tris, 150 mM sodium chloride, 5 mM DTT, pH 7.5, 25°C. This produced only minor changes in the HSQC spectrum of the unbound state so that assignments were readily transferred from above.

PRE measurements were made using ^15^N-labeled Cys1253Ala and Cys1253Ala/Thr1273Glu mutants of the S tail peptide that were reacted with 10 molar equivalents of MTSL (Santa Cruz Biotechnology) for 1-2 hours at room temperature. Reactions were checked for completion by MALDI and excess MTSL was removed by dialysis. Two-dimensional ^1^H-^15^N HSQC spectra were acquired on 100 μM MTSL-derived tail peptides before and after reduction with 20 molar equivalents of sodium ascorbate. Peak intensities for the oxidized and reduced states, *I*_ox_ and *I*_red_, were determined with SPARKY. To control for intermolecular effects, a natural abundance sample of the MTSL-peptide was mixed in a 1:1 ratio with a ^15^N-labeled Cys1253Ser/Cys1254Ser mutant S tail peptide, and peak intensities were measured for the oxidized and reduced states as above.

### Virus-like particles (VLPs)

HEK293T (obtained from Dr. Ed Campbell, Loyola University Chicago) and Vero-E6 (ATCC CRL-1586) cells were maintained in DMEM-10% FBS [Dulbecco’s Modified Eagle Media (DMEM) containing 10 mM HEPES, 100 nM sodium pyruvate, 0.1 mM non-essential amino acids, 100 U/ml penicillin G, and 100 μg/ml streptomycin, and supplemented with 10% fetal bovine serum (FBS, Atlanta Biologicals)]. These cell lines were cultured in a 5% CO2 incubator at 37°C. Full-length SARS-CoV S (GenBank: AY278741.1) and SARS-CoV-2 S, E, M, and N (GenBank: NC_045512.2) genes were synthesized by Genscript, Inc. as human codon-optimized cDNAs, and inserted into pcDNA3.1 expression vectors. HiBiT-N was constructed by fusing HiBiT peptide (VSGWRLFKKIS) coding sequences with linker (GSSGGSSG) to the 5’ end of the N gene, as described ^47,48,63^. The pCMV-LgBiT expression plasmid was purchased from Promega. pcDNA3.1-hACE2-C9 was obtained from Dr. Michael Farzan, Scripps Florida. pcDNA3.1-hACE2-LgBiT was constructed by fusing the coding sequence of LgBiT to the 3’ end of hACE2 gene. HiBiT-N tagged VLPs were produced as described previously ^47,48,63^. Briefly, equimolar amounts of full-length CoV S, E, M and HiBiT-N encoding plasmids (total 10 ug) were LipoD (SignaGen, cat: SL100668)-transfected into 107 HEK293T cells. To produce spike-less “No S” VLPs, the S expression plasmids were replaced with empty vector plasmids. At 6 h post-transfection, cells were replenished with fresh DMEM-10% FBS. HiBiT-N VLPs were collected in FBS-free DMEM from 24 to 48 h post-transfection. FBS-free DMEM containing HiBiT-N VLPs were clarified by centrifugation (300xg, 4°C, 10 min; 3000xg, 4°C, 10 min). To obtain purified viral particles, clarified VLP-containing FBS-free DMEM was concentrated 100-fold by ultracentrifugation (SW28, 7500 rpm, 4C, 24 h) through 20% (w/w) sucrose ^64^. VLPs were quantified after detergent lysis by adding LgBiT and measuring complemented Nluc in a luminometer. For downstream experiments, VLP inputs were normalized based on their Nluc activity upon LgBiT complementation. VLP samples were stored at -80°C.

### Cell-free fusion assay

hACE2-LgBiT extracellular vesicles (EVs) were obtained as described previously ^47,63^. Briefly, HEK293T target cells were LipoD-transfected with pcDNA3.1-hACE2-LgBiT. At 6 h post-transfection, transfection media were removed, rinsed, and replaced with FBS-free DMEM. Media were collected at 48 hours post-transfection, clarified (300xg, 4°C, 10 min; 3000xg, 4°C, 10 min), and concentrated 100-fold by ultrafiltration (Amicon, 100 kDa). EVs were then purified using size-exclusion chromatography (qEV original, Izon, Inc.) using PBS pH 7.4 as eluant. Peak EV fractions were identified by adding HiBiT-containing detergent and subsequent nano-luciferase (Nluc) measurement by luminometry. EVs were stored at 4°C. Cell-free fusion assays were performed as described previously (Qing et al., 2021; Qing et al., 2022). Briefly, at 4°C, equal volumes of HiBiT-N VLPs and hACE2-LgBiT EVs were mixed with nanoluc substrate (cat#N2420, Promega) and trypsin (Sigma; 10 ng/μl or as indicated) in 384-well multiwell plates. Sample plates were then loaded into a Glomax luminometer maintained at 37°C. Nluc accumulations were recorded over time. VLP-EV cell-free fusions were quantified as the fold increase of Nluc signal from S-bearing VLPs over the signal from spike-less (no S) VLP background control.

### Cell-cell fusion assay

VLP-producer cells and target cells were prepared as described previously ^47^. Briefly, VLP-producer HEK293T cells were co-transfected with pDSP1-7 and expression plasmids for SARS-CoV-2 S, E, M, and N-HiBiT. Control effector cells received empty vector plasmid instead of S plasmid. Target cells (HEK293T) were co-transfected with pDSP8-11 and empty vector or hACE2-expressing plasmids. At 6 hours post-transfection, VLP-producer and target cells were suspended and mixed into white-walled 96-well plates. 18 hours later, live-cell renilla luciferase (Rluc) substrate (EnduRen, Promega) was added. After 2 hours, Rluc levels were quantified using a Veritas microplate luminometer.

### Bottom-up proteomics

Samples were spiked with 10x lysis buffer (10 uL, 120 mM sodium lauroyl sarcosine, 5% sodium deoxycholate, 500 mM triethylammonium bicarbonate (TEAB), Halt™ Protease and Phosphatase Inhibitor Cocktail (Thermo Fisher Scientific)). The samples were treated with tris (2-carboxyethyl) phosphine (10 uL, 55 mM in 50 mM TEAB, 30 min, 60° C) followed by treatment with chloroacetamide (10 uL, 120 mM in 50 mM TEAB, 30 min, 25° C in the dark). They were then diluted 5-fold with aqueous 50 mM TEAB and incubated overnight with Sequencing Grade Modified Trypsin (1 ug in 10 uL of 50 mM TEAB; Promega, Madison, WI). Following this an equal volume of ethyl acetate/trifluoroacetic acid (TFA, 100/1, v/v) was added and after vigorous mixing (5 min) and centrifugation (13,000 x g, 5 min), the supernatants were discarded, and the lower phases were dried in a centrifugal vacuum concentrator. The samples were then desalted using a modified version of Rappsilber’s protocol ^65^ in which the dried samples were reconstituted in acetonitrile/water/TFA (solvent A, 100 uL, 2/98/0.1, v/v/v) and then loaded onto a small portion of a C18-silica disk (3M, Maplewood, MN) placed in a 200 uL pipette tip. Prior to sample loading the C18 disk was prepared by sequential treatment with methanol (20 uL), acetonitrile/water/TFA (solvent B, 20 uL, 80/20/0.1, v/v/v) and finally with solvent A (20 uL). After loading the sample, the disc was washed with solvent A (20 uL, eluent discarded) and eluted with solvent B (40 uL). The collected eluent was dried in a centrifugal vacuum concentrator and reconstituted in water/acetonitrile/FA (solvent E, 10 uL, 98/2/0.1, v/v/v). Aliquots (5 uL) were injected onto a reverse phase nanobore HPLC column (AcuTech Scientific, C18, 1.8um particle size, 360 um x 20 cm, 150 um ID), equilibrated in solvent E and eluted (5 00 nL/min) with an increasing concentration of solvent F (acetonitrile/water/FA, 98/2/0.1, v/v/v: min/% F; 0/0, 5/3, 18/7, 74/12, 144/24, 153/27, 162/40, 164/80, 174/80, 176/0, 180/0) using an EASY-nLC II (Thermo Fisher Scientific). The effluent from the column was directed to a nanospray ionization source connected to a hybrid quadrupole-Orbitrap mass spectrometer (Q Exactive Plus, Thermo Fisher Scientific) acquiring mass spectra in a data-dependent mode alternating between a full scan (m/z 350-1700, automated gain control (AGC) target 3 × 106, 50 ms maximum injection time, FWHM resolution 70,000 at m/z 200) and up to 15 MS/MS scans (quadrupole isolation of charge states 2-7, isolation window 0.7 m/z) with previously optimized fragmentation conditions (normalized collision energy of 32, dynamic exclusion of 30 s, AGC target 1 × 105, 100 ms maximum injection time, FWHM resolution 35,000 at m/z 200). The raw data was analyzed in Proteome Discoverer (Version 2.4; Thermo Fisher Scientific), which provided measurements of abundance for the identified peptides. Tryptic peptides containing amino acid sequences unique to individual proteins were used to identify coatomer subunits α, β, β’, o, ε, γ-1 and ζ-1 in each sample.

