## Supplementary Table 1 for "A single C-terminal residue controls SARS-CoV-2 spike trafficking and virion assembly"

**Extended Data Table S1:** Identification of coatomer subunits in pull-downs with clientized S tail

| **Accession** | **Gene Symbol** | **Description** | **Sum PEP Score** | **Sequence Coverage [%]** | **# Peptides** | **Abundance: Thr1273Asp** | **Abundance: Thr1273Glu** |
| --- | --- | --- | --- | --- | --- | --- | --- |
| Q8CIE6 | Copa | Coatomer subunit α | 126.501 | 23 | 22 | High | High |
| Q9JIF7 | Copb1 | Coatomer subunit β | 36.418 | 10 | 7 | High | High |
| O55029 | Copb2 | Coatomer subunit β' | 69.231 | 20 | 13 | High | High |
| Q5XJY5 | Arcn1 | Coatomer subunit δ | 1.588 | 2 | 1 | Peak Found | High |
| O89079 | Cope | Coatomer subunit ε | 31.878 | 34 | 7 | High | High |
| Q9QZE5 | Copg1 | Coatomer subunit γ-1 | 12.841 | 4 | 3 | High | High |
| P61924 | Copz1 | Coatomer subunit ζ-1 | 18.131 | 21 | 2 | High | High |
