## Supplementary Table 2 for "A single C-terminal residue controls SARS-CoV-2 spike trafficking and virion assembly"

**Extended Data Table S2:** Crystallographic data collection and refinement statistics.

| **Protein** | β'WD40 | β'WD40 | β'WD40-Tyr33Ala | αWD40-Arg13Ala | αWD40-Lys15Ala | αWD40-Arg300Ala |
| --- | --- | --- | --- | --- | --- | --- |
| S peptide | GVKLHYT | GVKLHYE | GVKLHYE | -- | -- | -- |
| PDB ID | 8ENS | 8ENW | 8ENX | 8ENY | 8ENZ | 8EO0 |
| **Data Collection^1^** | | | | | | |
| Space group | C121 | P1 | P1 | P1211 | P1211 | P1211 |
| Unit cell (a, b, c; Å) | 75.0, 51.4, 85.3 | 42.7, 46.0, 84.3 | 42.8, 46.2, 84.5 | 37.4, 170.2, 71.5 | 37.5, 172.3, 71.5 | 37.9, 171.3, 71.6 |
| Unit cell (α, β, γ; °) | 90.0, 101.7, 90.0 | 81.2, 81.5, 69.0 | 81.1, 81.7, 69.5 | 90.0, 99.8, 90.0 | 90.1, 99.9, 90.0 | 89.9, 100.3, 90.1 |
| Resolution (Å) | 28.1-1.4 (1.47-1.45) | 82.9-1.4 (1.47-1.45) | 28.3-1.8 (1.84-1.80) | 28.8-1.9 (1.94-1.90) | 28.8-1.6 (1.68-1.65) | 28.9-1.8 (1.83-1.80) |
| R_sym_ (%) | 10.7 (67.1) | 9.6 (88.4) | 13.7 (71.3) | 12.9 (84.5) | 7.0 (44.5) | 18.5 (19.6) |
| <I/σ(I)> | 9.1 (2.3) | 6.8 (1.4) | 3.7 (1.0) | 5.9 (1.4) | 8.4 (2.2) | 4.6 (1.1) |
| CC1/2 | 1.00 (0.84) | 1.00 (0.62) | 0.99 (0.55) | 0.99(0.57) | 0.99 (0.80) | 0.99 (0.55) |
| Redundancy | 6.9 (6.3) | 3.7 (3.8) | 2.4 (2.4) | 4.8 (4.4) | 3.6 (3.7) | 4.7 (4.7) |
| Total number of reflections | 390433 (17128) | 370084 (18804) | 128752 (7007) | 328403 (20135) | 381341 (19084) | 391365 (21430) |
| Number of unique reflections | 56398 (2705) | 100965 (4966) | 53029 (2959) | 67929 (4533) | 105242 (5174) | 82865 (4564) |
| Completeness (%) | 99.8 (96.4) | 96.6 (94.8) | 96.6 (90.2) | 98.2 (97.1) | 98.6 (98.2) | 100.0 (99.9) |
| Wilson B factor (Å^2^) | 14.4 | 15.2 | 16.1 | 23.9 | 17.7 | 20.2 |
| **Refinement** | | | | | | |
| Resolution used in refinement (Å) | 28.1-1.4 (1.47-1.45) | 42.7-1.4 (1.47-1.45) | 28.3-1.8 (1.84-1.80) | 28.4 - 1.9 (1.96 - 1.90) | 28.7 - 1.6 (1.71 - 1.65) | 28.9 - 1.8 (1.86 - 1.80) |
| Number of WD40 chains in ASU | 1 | 2 | 2 | 3 | 3 | 3 |
| Number of spike peptide chains in ASU | 1 | 2 | 1 | -- | -- | -- |
| Number of reflections used in refinement | 56384 (2736) | 100938 (3284) | 53018 (5144) | 67828 (6755) | 105146 (10362) | 82725 (8247) |
| R_work_ (%) | 15.1 (22.5) | 15.3 (25.6) | 14.9 (24.4) | 17.8 (25.1) | 16.3 (24.8) | 17.7 (28.3) |
| R_free_ (%)^2^ | 17.2 (24.0) | 18.0 (29.8) | 19.6 (28.8) | 21.9 (29.4) | 19.3 (26.4) | 21.8 (32.7) |
| Amino acid residues in final model | 308 | 617 | 611 | 917 | 917 | 915 |
| Atoms in final model | 5307 | 10385 | 10502 | 15162 | 15450 | 15177 |
| Water molecules | 469 | 530 | 826 | 453 | 735 | 525 |
| Hydrogen atoms | 2355 | 4839 | 4774 | 7234 | 7234 | 7203 |
| All atom clash-score | 1.6 | 1.8 | 2.0 | 2.9 | 2.9 | 2.7 |
| Ramachandran angles (favored, %) | 94.7 | 96.6 | 95.0 | 95.4 | 95.5 | 95.7 |
| Ramachandran angles (allowed, %) | 5.3 | 4.4 | 5.0 | 4.6 | 4.5 | 4.3 |
| Ramachandran angles (outliers, %) | 0.0 | 0.0 | 0.0 | 0.0 | 0.0 | 0.0 |
| Rotamers (outliers, %) | 0.7 | 0.0 | 0.0 | 0.2 | 0.74 | 0.4 |
| Cβ outliers | 0 | 0 | 0 | 0.0 | 0.0 | 0.0 |
| RMS bond length (Å) | 0.008 | 0.005 | 0.005 | 0.002 | 0.010 | 0.004 |
| RMS bond angle (°) | 1.086 | 0.946 | 0.759 | 0.600 | 1.030 | 0.770 |

^1^Data collected at NSLSII 17-ID-1 AMX beamline at an X-ray wavelength of 0.92Å. ^2^5% of reflections assigned for R_free_ calculations.
