## Supplementary Table 3 for "A single C-terminal residue controls SARS-CoV-2 spike trafficking and virion assembly"

| **Spike C-tail (1255-1273)** | **Conformation #** | **Number of tail atoms closer than 1Å to WD40 domain atoms** | **Number of tail atoms closer than 0.4Å to WD40 domain atoms** |
| --- | --- | --- | --- |
| WT | 1 | 3 | 0 |
|  | 2 | 4 | 1 |
|  | 3 | 10 | 2 |
|  | 4 | 7 | 0 |
|  | 5 | 4 | 0 |
|  | 6 | 2 | 0 |
|  | 7 | 2 | 0 |
|  | 8 | 6 | 0 |
|  | 9 | 4 | 0 |
|  | 10 | 3 | 1 |
|  | 11 | 5 | 0 |
|  | 12 | 4 | 0 |
|  | 13 | 11 | 1 |
|  | 14 | 5 | 1 |
|  | 15 | 21 | 2 |
|  | 16 | 8 | 4 |
|  | 17 | 2 | 0 |
|  | 18 | 14 | 2 |
|  | 19 | 0 | 0 |
|  | 20 | 0 | 0 |
| Thr1273Glu | 1 | 5 | 1 |
|  | 2 | 5 | 0 |
|  | 3 | 3 | 0 |
|  | 4 | 2 | 1 |
|  | 5 | 4 | 2 |
|  | 6 | 5 | 1 |
|  | 7 | 4 | 2 |
|  | 8 | 0 | 0 |
|  | 9 | 9 | 3 |
|  | 10 | 7 | 0 |
|  | 11 | 4 | 0 |
|  | 12 | 2 | 0 |
|  | 13 | 5 | 0 |
|  | 14 | 13 | 1 |
|  | 15 | 4 | 1 |
|  | 16 | 5 | 0 |
|  | 17 | 11 | 1 |
|  | 18 | 2 | 1 |
|  | 19 | 8 | 1 |
|  | 20 | 8 | 0 |

**Extended Data Table S3:** Structural analysis of wild-type and humanized S tail NMR ensembles for clashes with WD40 domain
