## Supplementary Figure S1 for "A single C-terminal residue controls SARS-CoV-2 spike trafficking and virion assembly"

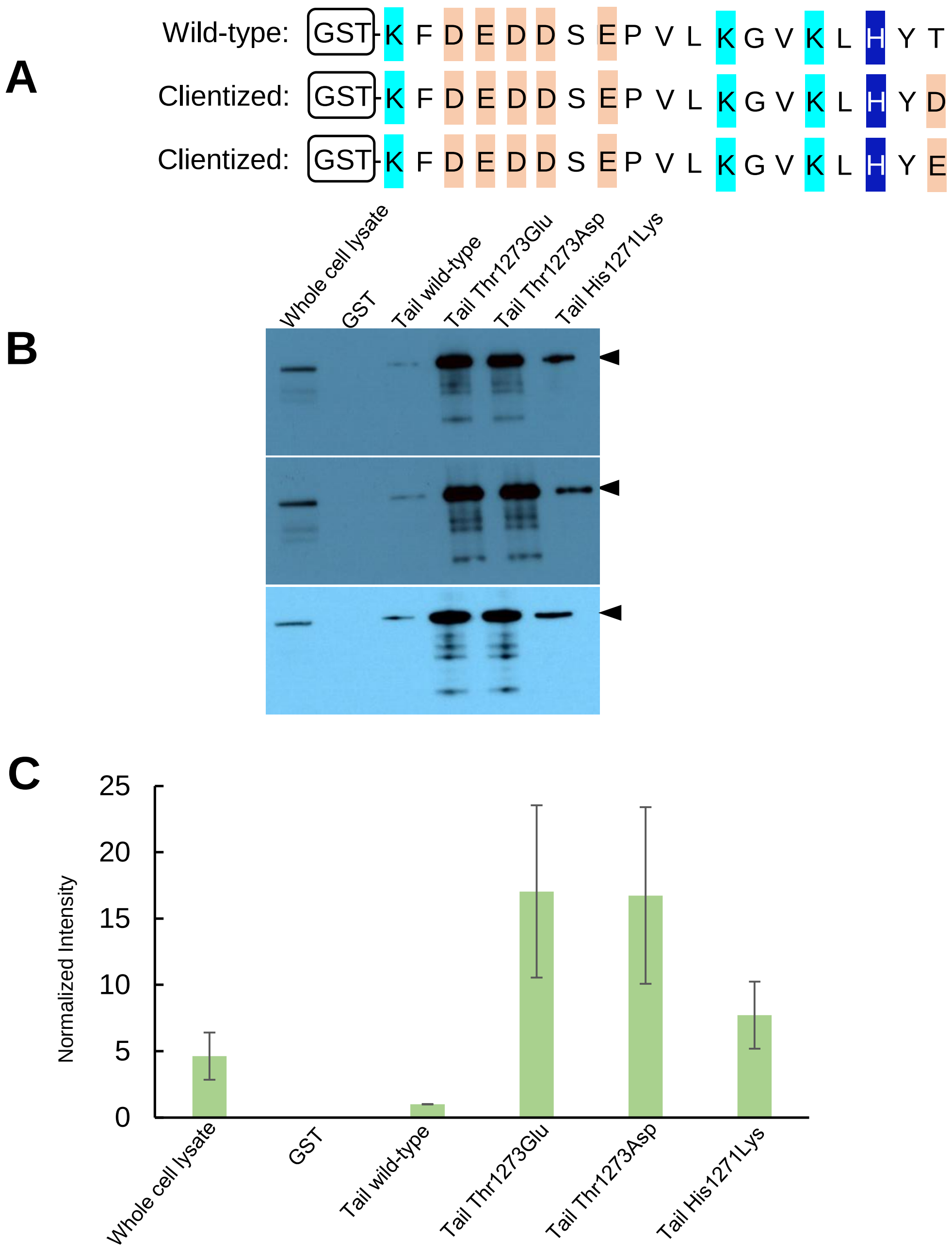


**Extended Data Figure S1.** Western blot analysis of coatomer pull-down by GST-S tail fusion protein. (A) Wild-type and clientized GST-tail constructs for coatomer pull-downs. (B) These panels show three technical replicates of pull-downs probed with an antibody for βCOPI subunit. (C) A bar-graph representation of mean intensity for the βCOPI subunit band. The data have been normalized to the intensity for the wild-type tail pull-down. Data represented as mean of triplicate independent samples ± SD.
