## Supplementary Figure S2 for "A single C-terminal residue controls SARS-CoV-2 spike trafficking and virion assembly"

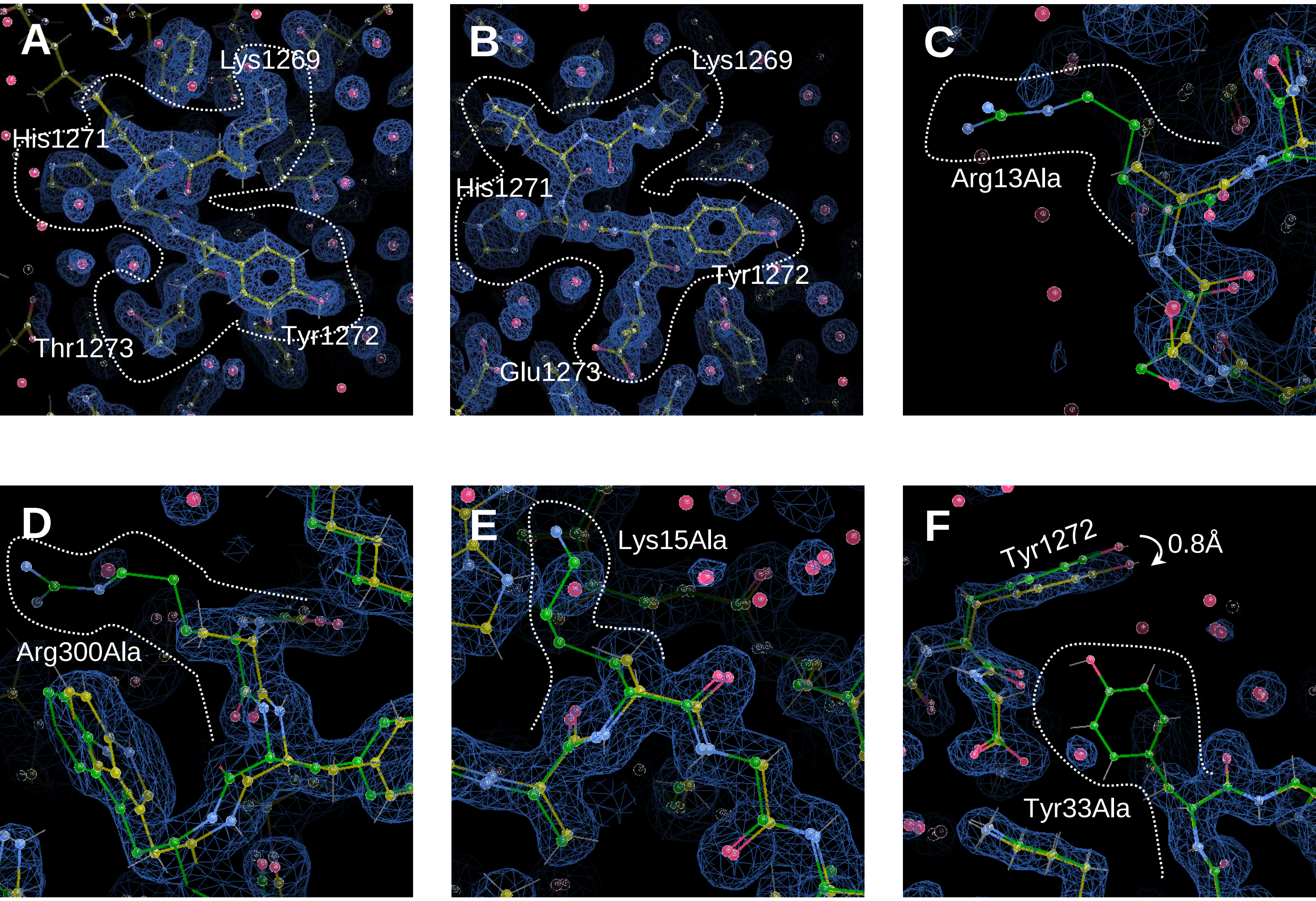


**Extended Data Figure S2.** Electron density of critical residues in S tail-WD40 interface highlighted with broken white lines. (A) Wild-type S tail hepta-peptide, (B) clientized S tail hepta-peptide, and the mutation sites at (C) Arg13Ala, (D) Arg300Ala, and (E) Lys15Ala. Coordinates corresponding to each electron density map are in yellow whereas superposed wild-type αWD40 in panels (C), (D), and (E) are in green. (F) The co-crystal structure of S tail hepta-peptide with wild-type β'WD40 (green) was superposed onto the co-crystal structure of S tail hepta-peptide with S tail hepta-peptide with β'WD40 Tyr33Ala mutant. The electron density map corresponds to the latter complex. Blue mesh: 2Fo-Fc map, σ=1.2.
