## Supplementary Figure S3 for "A single C-terminal residue controls SARS-CoV-2 spike trafficking and virion assembly"

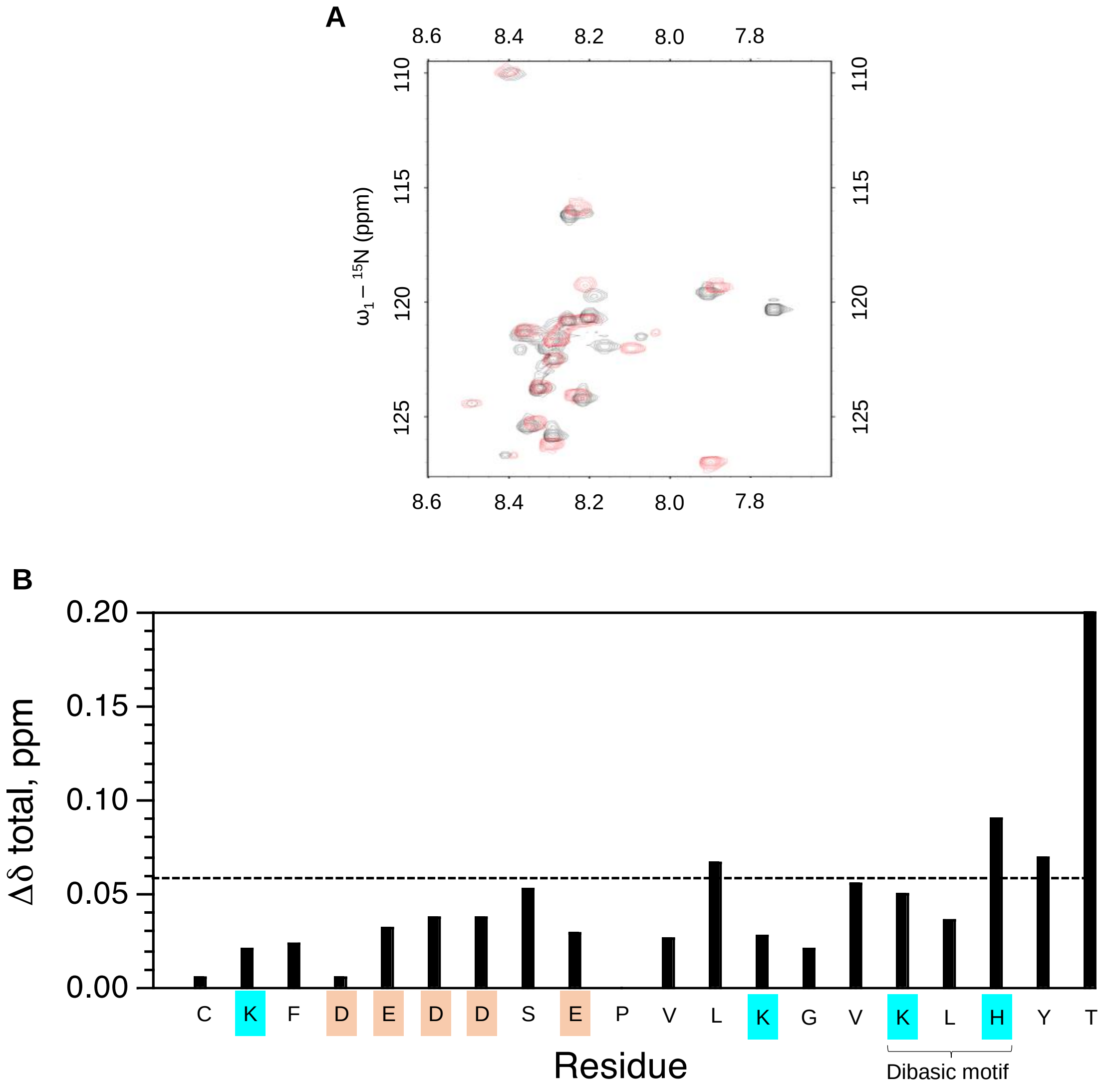


**Extended Data Figure S3.** (A) Overlaid 2D 1H-15N HSQC spectra of Cys1253Ala-21mer (black) and Cys1253Ala/Thr1273Glu-21mer (red). Both spectra were acquired at 25°C in 100 mM potassium phosphate , 0.1 mM DTT, pH 7.0. Peaks with notable shifts between the two spectra are labeled. (B) Plot of backbone amide chemical shift perturbations between the HSQC spectra for wild-type and clientized spike tails. Note that the plot is truncated as the Δ$\delta$ total value for Thr1273 is 1.35 ppm (asterisk). The dashed line indicates the mean value + 1SD, not including Thr1273.
