## Supplementary Figure S4 for "A single C-terminal residue controls SARS-CoV-2 spike trafficking and virion assembly"

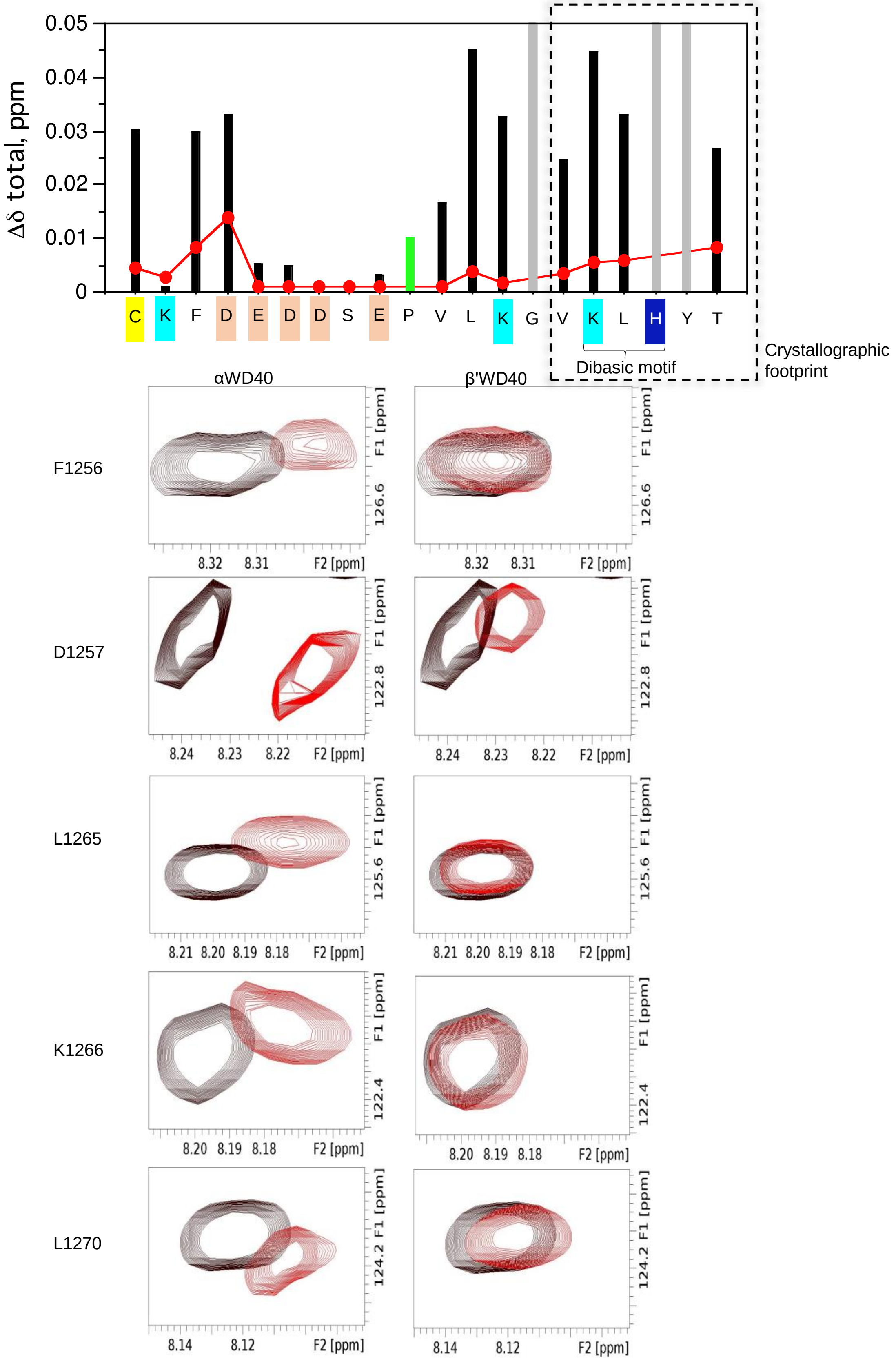


**Extended Data Figure S4.** Larger S tail CSPs observed with αWD40 than β'WD40. Upper panel: Backbone amide CSP plots for wild-type $^{15}$N-S tail 21mer peptide upon addition of unlabeled αWD40 (black) or β'WD40 (red); gray bars indicate exchange broadened peaks with addition of either αWD40 or β'WD40; green bar, Pro1263 (no CSP). Lower panel: Regions from the two dimensional $^{1}$H-$^{15}$N HSQC spectrum of wild-type 15N-S tail 21mer peptide upon addition of 1:1 equivalent of either unlabeled αWD40 (left) or unlabeled β'WD40 (right); free peptide (black); complex (red).
