## Supplementary Figure S5 for "A single C-terminal residue controls SARS-CoV-2 spike trafficking and virion assembly"

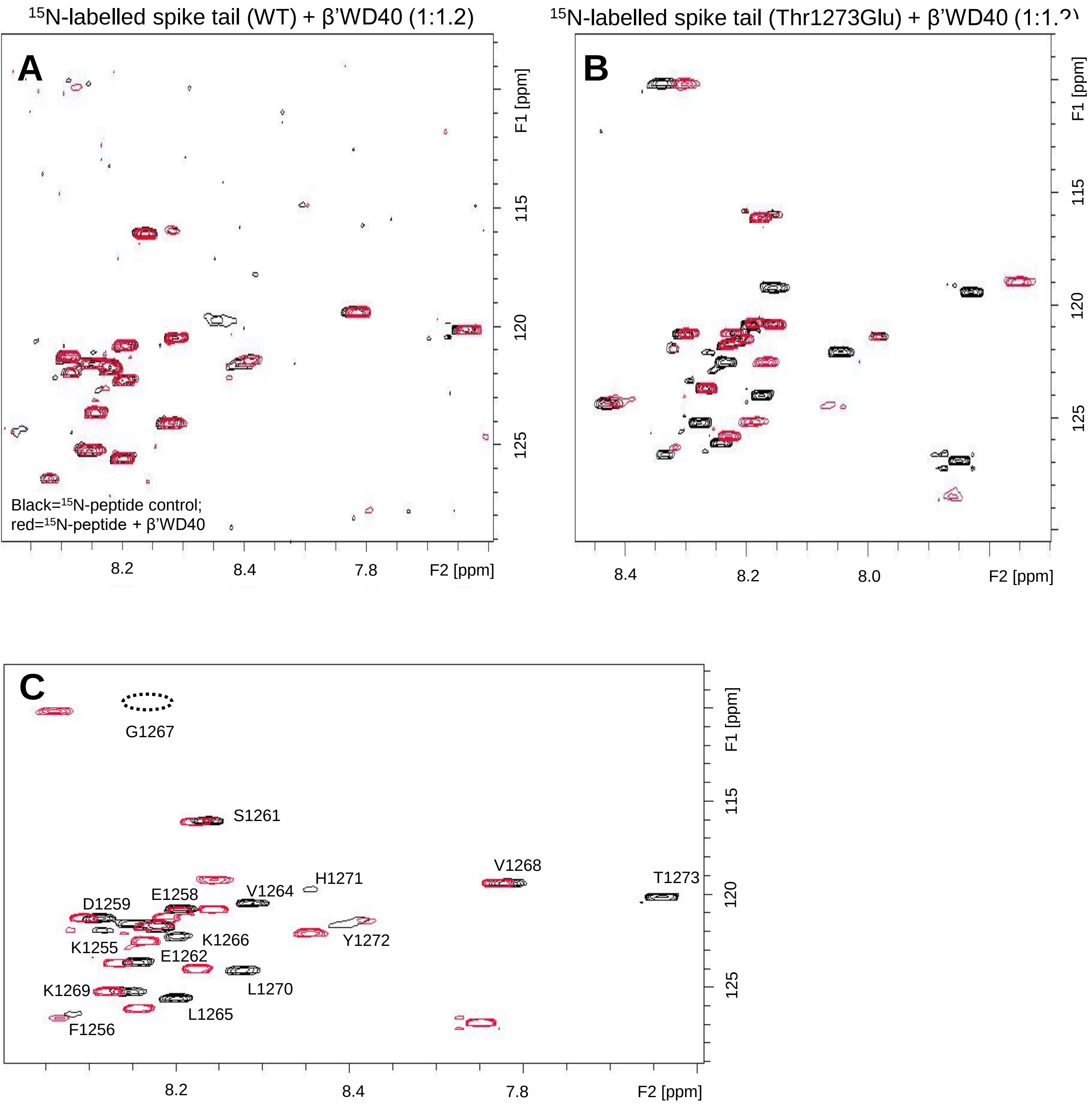


**Extended Data Figure S5.** HSQC analysis shows, (A) weak interaction of β’WD40 with the wild-type spike tail but, (B) substantially stronger interaction with the clientized tail as inferred from large CSPs. (C) This panel shows residue assignments for the wild-type spike tail.
