## Supplementary Figure S6 for "A single C-terminal residue controls SARS-CoV-2 spike trafficking and virion assembly"

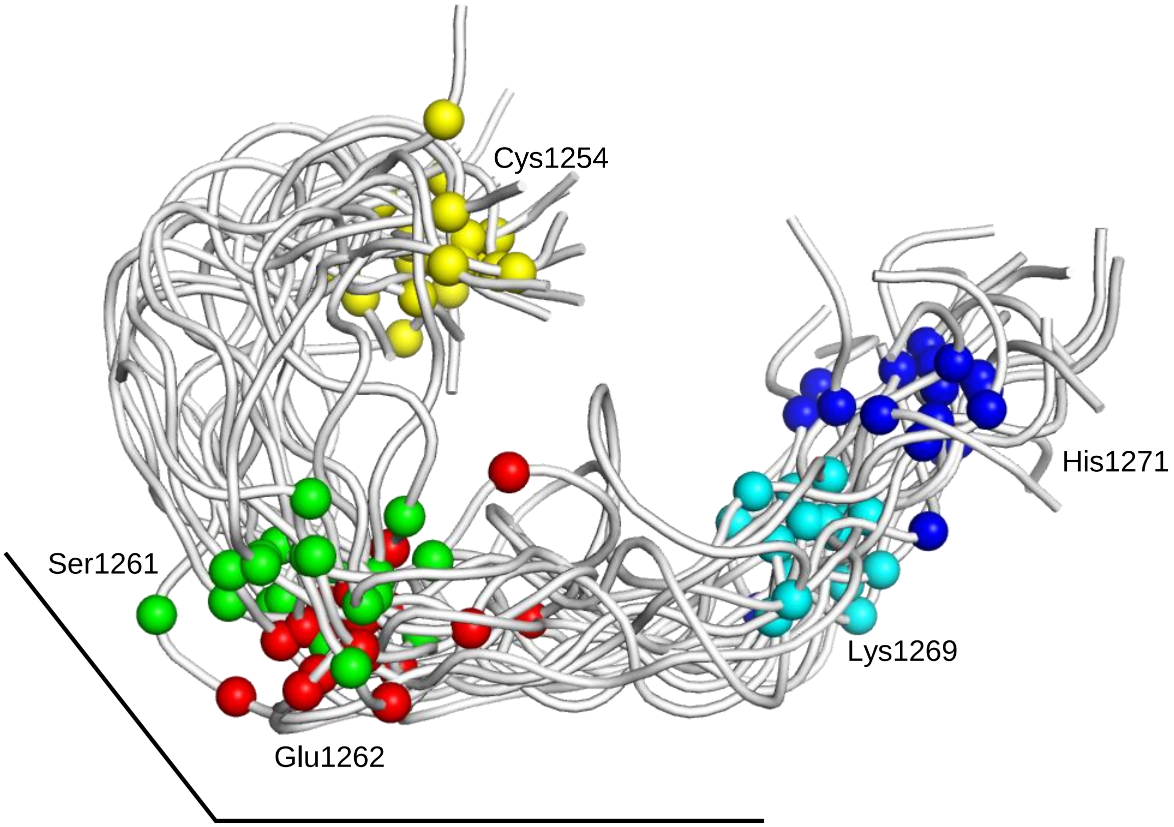


**Extended Data Figure S6.** Ensemble of wild-type S tail showing Cα atoms for Cys1254 (yellow), Ser1261 (green), Glu1262 (red), Lys1269 (cyan), and His1272 (blue). The black lines highlight the turn proximal to Ser1261-Glu1262 in the S tail main-chain.
